# Predictive nonlinear modeling of malignant myelopoiesis and tyrosine kinase inhibitor therapy

**DOI:** 10.1101/2022.10.11.511822

**Authors:** Jonathan Rodriguez, Abdon Iniguez, Nilamani Jena, Prasanthi Tata, Joan Liu, Arthur D. Lander, John S. Lowengrub, Richard A. Van Etten

**Affiliations:** Graduate Program in Mathematical, Computational & Systems Biology, University of California, Irvine, Irvine CA 92697; Center for Complex Biological Systems, University of California, Irvine, Irvine CA 92697; Department of Medicine, University of California, Irvine, Irvine CA 92697; Department of Developmental and Cell Biology, University of California, Irvine, Irvine CA 92697; Chao Family Comprehensive Cancer Center, University of California, Irvine, Irvine CA 92697; Department of Mathematics, University of California, Irvine, Irvine CA 92697; Department of Biomedical Engineering, University of California, Irvine, Irvine CA 92697

## Abstract

Chronic myeloid leukemia (CML) is a blood cancer characterized by dysregulated production of maturing myeloid cells driven by the product of the Philadelphia chromosome, the BCR-ABL1 tyrosine kinase. Tyrosine kinase inhibitors (TKI) have proved effective in treating CML but there is still a cohort of patients who do not respond to TKI therapy even in the absence of mutations in the BCR-ABL1 kinase domain that mediate drug resistance. To discover novel strategies to improve TKI therapy in CML, we developed a nonlinear mathematical model of CML hematopoiesis that incorporates feedback control and lineage branching. Cell-cell interactions were constrained using an automated model selection method together with previous observations and new *in vivo* data from a chimeric *BCR-ABL1* transgenic mouse model of CML. The resulting quantitative model captures the dynamics of normal and CML cells at various stages of the disease and exhibits variable responses to TKI treatment, consistent with those of CML patients. The model predicts that an increase in the proportion of CML stem cells in the bone marrow would decrease the tendency of the disease to respond to TKI therapy, in concordance with clinical data and confirmed experimentally in mice. The model further suggests that a key predictor of refractory response to TKI treatment is an increased probability of self-renewal of normal hematopoietic stem cells. We use these insights to develop a clinical prognostic criterion to predict the efficacy of TKI treatment and to design strategies to improve treatment response. The model predicts that stimulating the differentiation of leukemic stem cells while applying TKI therapy can significantly improve treatment outcomes.

## Introduction

Chronic myeloid leukemia (CML) is a myeloproliferative neoplasm of the hematopoietic system, which normally produces billions of mature myeloid and erythroid cells on a daily basis, is tightly regulated, and accommodates massive increases in production of individual cell types in response to physiologic and pathologic stresses. The hematopoietic system is organized hierarchically as a collection of progressively more differentiated cells starting from a hematopoietic stem cell (HSC) located in the bone marrow (BM) and ending with post-mitotic terminally differentiated myeloid and lymphoid cells (*Rieger and Schroeder, 2012; Liggett and Sankaran, 2020*).

CML is characterized by an overproduction of myeloid cells including mature granulocytes (neutrophils, basophils and eosinophils) and their immediate precursors (metamyelocytes, myelocytes, promyelocytes), and of myeloid progenitors (*Jamieson et al., 2004*) including multipotential progenitors (MPPs) and committed progenitors (common myeloid progenitors (CMP), granulocyte-macrophage progenitors (GMP), and megakaryocyte-erythroid progenitors (MEP)). Untreated, the disease has three distinct phases (*Chereda and Melo, 2015*). In the initial “chronic” phase, the differentiation of myeloid progenitors is essentially normal, resulting in excessive levels of mature post-mitotic neutrophils and their immediate precursors. In later stages of the disease (accelerated phase and blast crisis), differentiation is reduced and expansion of immature progenitors is observed. Additional clonal karyotypic abnormalities are typically only observed during the accelerated and blast crisis phases (*Hehlmann et al., 2020*).

CML has one of the simplest of cancer genomes. It is driven by a single genetic abnormality arising somatically in an HSC, the Philadelphia (Ph) chromosome, the result of a balanced translocation between chromosomes 9 and 22 that creates a fusion of the genes for *BCR* and *ABL1*. The product of the *BCR-ABL1* fusion gene is a dysregulated cytoplasmic protein-tyrosine kinase, BCR-ABL1. CML thus represents a natural model of dysregulated granulocytopoiesis (*Quintás-Cardama and Cortes, 2009*).

Cell biological studies have shown that Ph^+^ cells expressing markers of normal HSC are capable of engrafting immunodeficient mice (*Sirard et al., 1996; Lewis et al., 1998*), implying that these cells are leukemia-initiating or leukemic “stem” cells (LSC). More mature committed progenitors in CML, like normal progenitors, lack sustained self-renewal capacity and cannot stably engraft immunodeficient mice nor generate hematopoietic colonies *in vitro* upon serial replating (*Huntly et al., 2004*). The proportion of LSCs in the BM is highly variable across CML patients at diagnosis and can range from a few percent to nearly one hundred percent (*Petzer et al., 1996; Diaz-Blanco et al., 2007; Abe et al., 2008; Thielen et al., 2016*), perhaps reflecting different periods of time patients spend in chronic phase before they are diagnosed, different rates of disease progression, or both.

There is persuasive experimental evidence of significant feedback regulation of different cell compartments in the dynamics of myeloid cell production in both normal and CML hematopoiesis, including signaling between the normal and CML cells (*Jiang et al., 1999; Devireddy et al., 2005; Vicente-Duenas et al., 2009; Naka et al., 2010; Reynaud et al., 2011; Zhang et al., 2012; Krause et al., 2013; Walenda et al., 2014; Welner et al., 2015*). For instance, experiments in a mouse model of CML provided evidence that IL-6, produced by leukemic neutrophils, blocked MPP differentiation towards a lymphoid fate, implying feedback from the myeloid lineage onto MPPs (*Reynaud et al., 2011*). Surprisingly, our knowledge of the details of feedback regulation in hematopoiesis is still incomplete, especially for granulopoiesis, where even late-stage feedback interactions are poorly understood. For example, two cytokines, granulocyte colony-stimulating factor (G-CSF) and granulocyte-macrophage colony-stimulating factor (GM-CSF), can pharmacologically increase neutrophil production, but mice lacking both cytokines maintain baseline neutrophil levels and can still increase neutrophil production in response to infection (*Basu et al., 2000*). In many cases, it is not known which cell types are providing and receiving the feedback, what signals are used, and what aspects of proliferative cell behavior they influence (i.e., proliferation rates, renewal probability, or progeny fate choice).

In spite of this knowledge deficit, CML can be treated quite effectively using selective small-molecule tyrosine kinase inhibitors of the BCR-ABL1 kinase. Tyrosine kinase inhibitors (TKI) such as imatinib, dasatinib and nilotinib, which inhibit proliferation and increase apoptosis of Ph^+^ cells, have dramatically lowered CML death rates (*Gambacorti-Passerini et al., 2011. The response to TKI therapy in CML is monitored primarily by determining the level of BCR-ABL1 mRNA transcripts in peripheral blood, normalized to a control RNA and expressed as a percentage on an International Scale* {*Arora, 2017 #99*). *BCR-ABL1* transcript levels, an approximation of the proportion of circulating malignant cells at any given time, generally decrease exponentially in patients responding to TKI therapy resulting in at least two distinct slopes when plotted semi-logarithmically—an initial rapid decline attributed to TKI-induced killing of more mature myeloid cells, and a subsequent slower decline postulated to represent lower death rates of more primitive leukemic stem/progenitor cells (*Michor et al., 2005*). Clinical resistance to TKI therapy in CML is a significant problem, and is classified as acquired resistance (increasing *BCR-ABL1* transcript levels following a substantial decrease) or primary resistance (lack of an adequate initial response). Many patients with acquired resistance have developed mutations in the BCR-ABL1 kinase domain that mediate pharmacological resistance to the TKI (*Ernst and Hochhaus, 2012*). By contrast, 10-15% of newly diagnosed CML patients fail to achieve an “early molecular response”, defined as the level of *BCR-ABL1* transcripts being less than 10% at 3 months (*Hanfstein et al., 2012; Marin et al., 2012*). Clinical data indicate that switching TKIs may not benefit these patients (*Yeung et al., 2012; Yeung et al., 2015*), suggesting that this group is destined to do poorly regardless of the specific inhibitor used. *BCR-ABL1* mutations are generally not present in this group of patients (*Zhang et al., 2009; Pietarinen et al., 2016*), and thus the mechanism(s) underlying this primary resistance is unclear. We hypothesized that these variable patient responses to TKI therapy arise from nonlinearity introduced by non-cell autonomous interactions between normal and CML cells. To test this hypothesis, we developed a novel mathematical model of CML hematopoiesis and TKI treatment that incorporates lineage branching and interactions between normal and CML cells through feedback and feedforward regulation.

Mathematical modeling of leukemia has a long history aimed at understanding disease progression and improving treatment response using single and combination targeted therapies and immunotherapy (*Whichard et al., 2010; Pujo-Menjouet, 2015; Brunetti et al., 2021; Kuznetsov et al., 2021; Roeder and Glauche, 2021*). Further, recent efforts have been made to integrate mathematical modeling in clinical decision-making to design personalized therapies (*Hoffmann et al., 2020; Engelhardt and Michor, 2021*). Many models of leukemia have utilized simplified lineage architectures and minimal feedback (*Roeder et al., 2006; Komarova and Wodarz, 2007; Horn et al., 2008; Foo et al., 2009; Hähnel et al., 2020; Pedersen et al., 2021*). While these models can be made to fit the multiphasic disease response data of CML to TKI treatment, the simplicity of the models can make these fitted parameters of limited clinical value. More physiologically accurate, nonlinear models that account for cell-cell signaling and lineage branching is expected to improve clinical relevance. Mathematical models that incorporate feedback signaling have been developed in normal (*Engel et al., 2004; Marciniak-Czochra et al., 2009; Mahadik et al., 2019*) (*Mon Père et al., 2021*) and diseased (*Wodarz, 2008; Sachs et al., 2011; Krinner et al., 2013; Stiehl et al., 2014; Stiehl et al., 2015; Crowell et al., 2016; Woywod et al., 2017; Jiao et al., 2018; Stiehl et al., 2018; Zenati et al., 2018; Park et al., 2019; Sharp et al., 2020*) hematopoiesis. Because of the vast number of possible ways in which feedback models of normal hematopoiesis and leukemia can be configured, mathematical models tend to greatly simplify the lineage architectures and the feedback interactions among the cell types. For example, Manesso and colleagues(*Manesso et al., 2013*) developed a hierarchical ordinary differential equation (ODE) model of normal hematopoiesis containing multiple cell types and branch points (16 cell types and 4 branch points) in the lineage tree. Limiting the feedback loops to involve only local, negative regulation (e.g., regulation by self and immediate progenitor/progeny in the lineage tree) results in about 10^6^ models, which enabled the use of a stochastic optimization algorithm to obtain parameters consistent with homeostasis and a requirement for a rapid return to equilibrium following system perturbations.

In the context of leukemia, the model architectures are typically much simpler. Generally, models of leukemia introduce a parallel mutant lineage with the same structure as that used to model the normal hematopoietic cells but with different parameters. For example, Wodarz developed an unbranched lineage ordinary differential equation (ODE) model of normal and leukemia stem and differentiated cells in which feedback from the differentiated cells controlled whether the stem cells divided symmetrically or asymmetrically, and demonstrated this provides a mechanism for blast crisis in CML to occur without additional mutations (*Wodarz, 2008*). Krinner et al. incorporated positive and negative feedback regulation of differentiation and proliferation in an unbranched lineage model that combined a discrete agent-based model for the stem cell compartment with an ODE system for the progenitor and differentiated cells to provide a detailed view of the stem cell dynamics and to test the effect of therapies (*Krinner et al., 2013*). Stiehl and co-workers (*Stiehl et al., 2015*) developed an unbranched lineage ODE model of normal and leukemic cells in which only negative feedback regulation of stem and progenitor cell self-renewal fractions was considered and this was further limited to arise only from factors produced by the post-mitotic, mature normal and leukemic cells. Later work extended this approach to investigate clonal selection and therapy resistance (*Stiehl et al., 2014*), the role of cytokines on leukemia progression (*Stiehl et al., 2018*), combination treatment strategies (*Banck and Görlich, 2019*) and niche competition (*Stiehl et al., 2020*). Clonal competition was also considered in an ODE feedback model of CML (*Woywod et al., 2017*) and a stochastic model with feedback (*Dinh et al., 2021*). Simpler unbranched lineage models of normal and leukemic cells in which only the normal cells respond to feedback but normal and leukemic cells compete for space in the bone marrow have been used to investigate regimes of co-existence of normal and leukemic cells (*Crowell et al., 2016; Jiao et al., 2018*) and to design combination therapies using optimal control algorithms (*Sharp et al., 2020*).

Here, we develop a nonlinear ordinary differential equation (ODE) model of normal and CML hematopoiesis using a general approach that integrates an automated method, Design Space Analysis (DSA)(*Fasani and Savageau, 2010*) with data gleaned from previously published experiments, and from two new *in vivo* experiments presented here that separately decrement the number of stem cells and terminally-differentiated myeloid cells in the bone marrow of mice. This approach enables us to systematically select among plausible model architectures and signaling interactions without a priori knowledge of which cells are providing and receiving signaling stimuli. We start with a model for normal hematopoiesis that accounts for stem, multipotent progenitor cells, and two types of terminally-differentiated cells representing the myeloid and lymphoid lineage branches. This approach allows us to reduce the potential model space from about 60,000 models to a single model class and reveals the existence of feedforward and feedback mechanisms. We then extend the model to incorporate CML hematopoiesis by introducing a parallel lineage of CML cells with the same model architecture but with different parameters. The model captures the dynamics of CML at various stages of the disease and exhibits variable response to TKI treatment consistent with that observed in clinical data. The model suggests biomarkers of primary resistance, identifies the underlying mechanisms governing the response to TKI therapy, and suggests new treatment strategies.

## Results

### Model of normal hematopoiesis

The primary challenge in developing mathematical models of normal and CML hematopoiesis is sorting through the combinatorial explosion of models that occurs when cell-cell signaling interactions are taken into account. Consider the model hematopoietic system shown in **Fig. 1A**, which accounts for hematopoietic stem (**HSC**; **S**), multipotent progenitor (**MPP**; **P**) and two types of post-mitotic, terminally differentiated cells--myeloid (**TD**_**m**_) and lymphoid (**TD**_**l**_). The HSC self-renew with fraction (e.g., probability) p_0_ or differentiate with fraction 1-p_0_. That is, the fraction of HSC that remain as HSC after division is p_0_. The MPPs self-renew with fraction p_1_ and differentiate into either lymphoid or myeloid cells with fractions q_1_ and 1-p_1_-q_1_, respectively. The HSC and MPPs divide with rates η_1_ and η_2_ and the myeloid and lymphoid cells die at rates d_m_ and d_l_, respectively. The ordinary differential equations (ODEs) that govern the dynamics of the cells are given in Methods. Assuming that there is either positive or negative regulation of the self-renewal and differentiation probabilities and division rates of any cell type from any other cell type results in 59,049 models, counting each combination of regulated signaling as a separate model.

**Figure 1:**
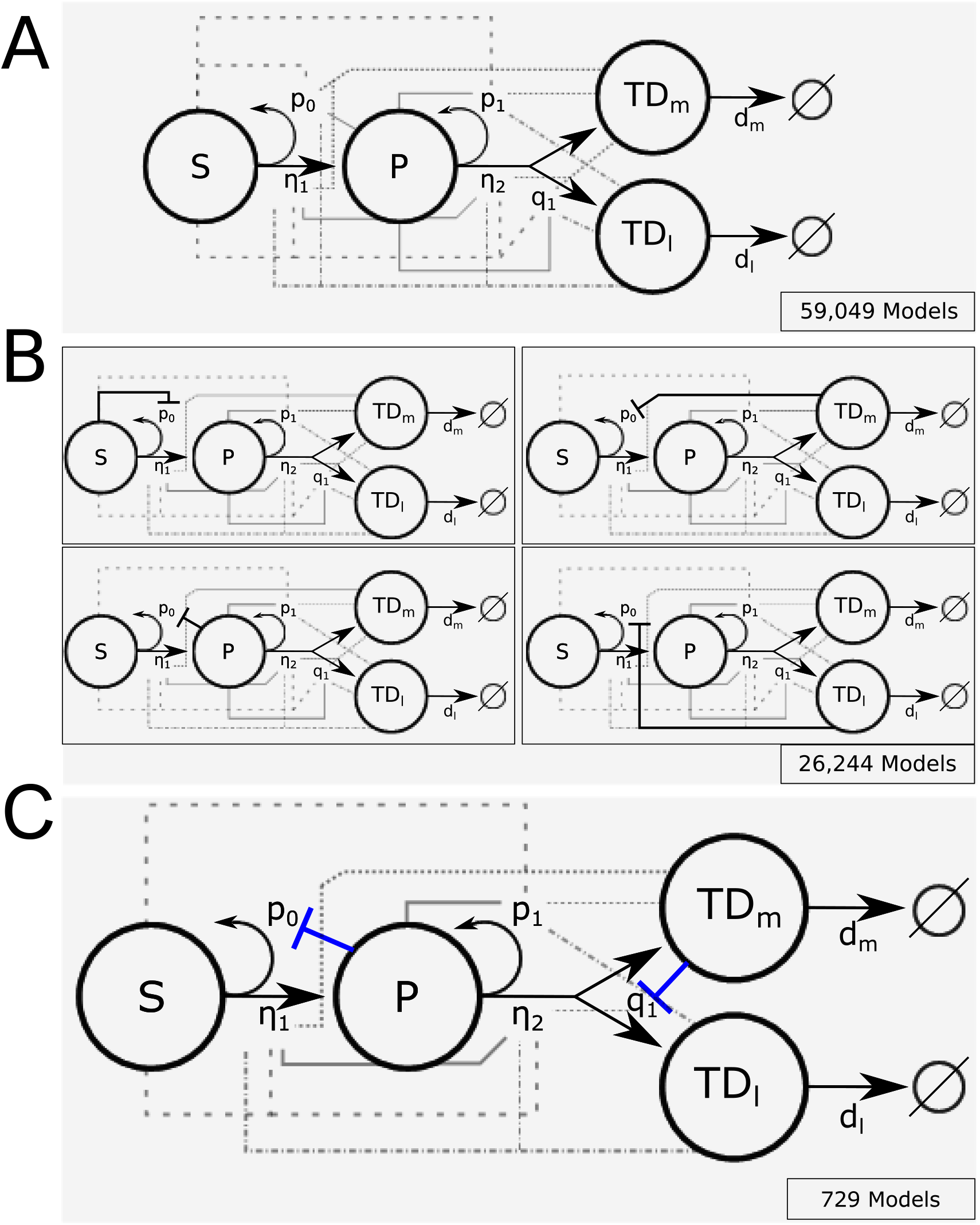
Branched lineage model of normal hematopoiesis with feedback regulation. **A**. Branched lineage model consisting of hematopoetic stem cells (HSC; S), multipotent progenitor cells (MPP; P), and post-mitotic, terminally differentiated myeloid (TD_m_) and lymphoid (TD_l_) cells. Modulation of the HSC and MPP self-renewal fractions (p_0_ and p_1_), division rates (η_1_ and η_2_), and fate switching probability (q_1_) through feedback can arise from any cell type. The different line styles correspond to regulation by a particular cell type (dashed for S, solid for P, dot-dashed for TD_l_, and dotted for TD_m_). **B**. Using Design Space Analysis, 4 candidate model classes are identified that differ in how HSCs are regulated. **C**. Using biological data from the literature as discussed in the text, we reduced the model space by hypothesizing that factors secreted by terminally differentiated myeloid cells direct the fate of MPPs (e.g., IL-6) and those by MPPs suppress HSC self-renewal (e.g., CCL3).

To select the most physiologically accurate models, we first filtered the models using an automated approach (Design Space Analysis or DSA) developed by Savageau and co-workers (*Savageau et al., 2009; Fasani and Savageau, 2010; Lomnitz and Savageau, 2016*) that enables models to be distinguished based on their range of qualitatively distinct behaviors, without relying on knowledge of specific values of the parameters. This method relies on identifying boundaries in parameter space that separate qualitative behaviors of a particular model, which is much more efficient than searching for model behaviors directly. The boundaries can be approximated from a sequence of inequalities that identify regions where one term on the right-hand side of each ODE (e.g., the rate of change) dominates all others in the sources (positive terms) and another dominates the sinks (negative terms). This is known as a dominant subsystem (S-system) of the model. The number of S-systems in each model depends on the number of combinations of positive and negative terms in the rates of change. If the equilibria of the S-systems, which are determined analytically, are not self-consistent (e.g., consistent with the assumed dominance of terms reflected in the inequalities) or the equilibria are not stable, then the S-system is rejected. If all the S-systems of a particular model are rejected, then that model is removed from further consideration. Models with at least one self-consistent and stable S-system are viable candidates for further analysis. DSA can be easily automated to make the analysis of very large numbers of equations feasible. Details are provided in Methods and the Supplemental Material (**Sec. S1**). The result of this procedure is the elimination of all but the 4 model classes shown in **Fig. 1B**, which require negative regulation of the stem cell self-renewal fraction but differ by where this regulation arises. The models within the classes share at least one S-system and have common qualitative behaviors. The differences between models in a class lie in whether or not there is positive, negative or no regulation on the rest of the parameters from any of the cell types. This reduces the number of possible models to 26,244.

Previous work has implicated several feedback mechanisms active in both normal and malignant hematopoiesis. Interleukin-6 (IL-6) is produced by differentiated myeloid cells and acts to bias MPPs towards a myeloid fate (*Reynaud et al., 2011; Welner et al., 2015*). Such negative feedback circuits, known as fate control, have been shown to provide an effective strategy for robust control of cell proliferation and reduction of oscillations in branched lineages (*Buzi et al., 2015*). The chemokine CCL3 (also known as macrophage inhibitory protein α or MIP-1α), produced in BM by basophilic myeloid progenitors (*Baba et al., 2016*), acts to inhibit the proliferation and self-renewal of normal HSC (*Broxmeyer et al., 1989; Staversky et al., 2018*), but CML HSC are relatively resistant to its action (*Eaves et al., 1993; Baba et al., 2013*). In hypothesizing these regulatory networks, we arrived at a single model class as shown in **Fig. 1C**. In this class, there are 729 model candidates, which differ only how the HSC and MPP cell division rates and the MPP self-renewal fraction are regulated. These above results suggest that IL-6 is a candidate feedback factor expressed in the myeloid compartment (TD_m_) with the ability to negatively regulate the fraction q_1_ of MPPs that differentiate into lymphoid cells. CCL3 is a candidate factor mediating negative feedback from the MPP population onto HSC self-renewal. To further constrain the remaining models, we performed cell biological experiments in mice to glean information about cell-cell interactions by separately perturbing the stem cell and myeloid cell compartments.

### Depletion of HSC increases HSC and MPP proliferation

As described in Methods, healthy C57BL6/J (B6) mice were treated with low-dose (50 cGy) ionizing radiation, previously shown to be selectively toxic to HSC in the bone marrow (BM) (*Stewart et al., 1998*). The BM stem/progenitor compartment was analyzed by flow cytometry in untreated mice, and on days 1, 3, and 7 post-irradiation, using the gating strategy in **Fig. 2A**. These time points and the number of mice analyzed at each time point were informed by a Bayesian hierarchical framework for optimal experimental design of mathematical models of hematopoeisis (*Lomeli et al., 2021*), One day after treatment, we observed an acute ∼2-fold decrease in the relative size of the HSC compartment in the irradiated mice (**Fig. 2B**), accompanied by ∼3-fold increase in proliferation rates for both HSC and MPPs (**Fig. 2C**). There was no change in MPP population, however, and the system returned to equilibrium by day 7. These results suggest that the HSC population exerts negative feedback on their own division rate (η_1_) and inhibits the division of MPPs through a negative feedforward loop on η_2_.

**Figure 2:**
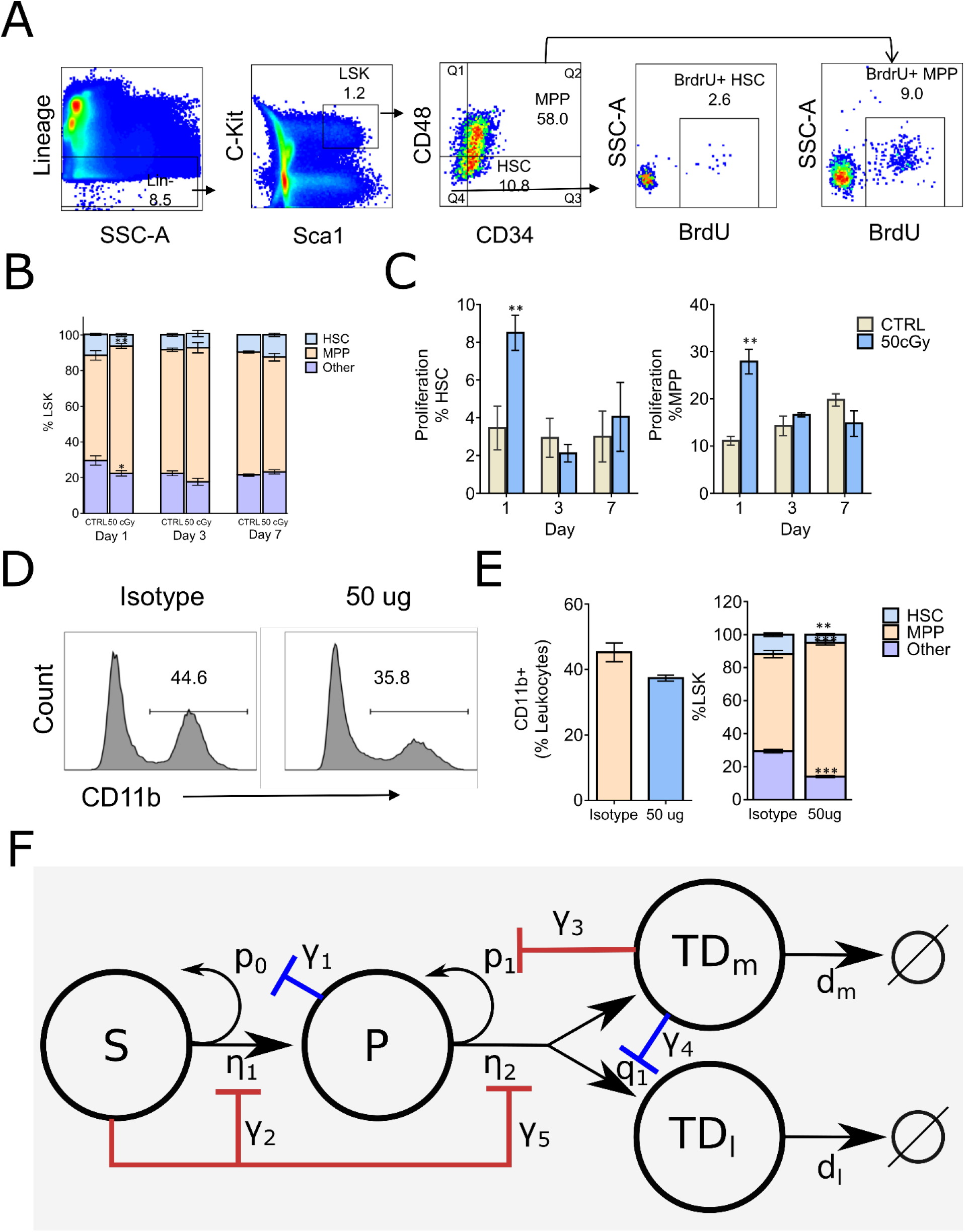
Fluorescence-activated cell sorting (FACS) analysis of mouse Lin^−^Sca-1^+^c-Kit^+^ (LSK) bone marrow stem/progenitor cells and the proposed branched lineage hematopoiesis model. **A**. Gating schema for phenotyping Hematopoietic Stem Cells (HSC, defined as LSK CD34^−^ CD48^−^) and Multi-Potential Progenitors (MPP, defined as LSK CD34^+^CD48^+^), and BrdU incorporation in their respective compartments. **B**. Distributions of HSC (blue), MPP (orange), and other (purple) compartments on days 1, 3, and 7 in the bone marrow of control (CTRL) B6 mice and mice that received 50 cGy radiation. **C**. Frequency of HSC and MPP proliferation in CTRL (gray bars) and irradiated (blue bars) mice measured by BrdU incorporation on days 1, 3, and 7. Data are shown as mean ± SEM. *p < 0.05. **D**. Representative histograms depicting the frequency of myeloid cells as measured by CD11b expression in mice 24 hrs after intravenous administration of isotype control (iso) or RB68C5 (50 μg) antibody. **E**. Left panel: Bar graph showing the frequency of CD11b^+^ cells in BM of mice that were treated with isotype control antibody (Iso; orange bar, n=3) or RB68C5 antibody (50 μg; blue bars, n=3). Right panel: HSC (blue), MPP (orange), and other cell type (purple) frequencies from mice that received isotype or RB6-8C5 antibody. Data are shown as mean ± SEM. *p < 0.05 **F**. Proposed feedforward-feedback model of hematopoiesis. The negative feedback loops shown in blue correspond to those suggested by previous experimental data (*Reynaud et al., 2011; Staversky et al., 2018*), while the negative feedback and feedforward loops in red are supported by our cell depletion experiments in **A-E**.

### Depletion of mature myeloid cells increases the MPP population

B6 mice were treated with the anti-granulocyte antibody RB68C5 (50 μg) and their BM was analyzed one day after treatment (see Methods). This treatment resulted in a ∼20% decrease in mature BM myeloid cells, as measured by CD11b expression (**Figs. 2D** and **E**), and was accompanied by a concomitant increase in the size of the phenotypic MPP compartment (**Fig. 2E**) and a decrease in the HSC compartment (**Fig. 2E**). These results suggest that there is a negative feedback loop from the myeloid cells onto the MPP self-renewal fraction p_1_.

Taking all these results into consideration, we arrive at the feedback-feedforward model shown in **Fig. 2F**. The negative feedback loops shown in blue correspond to those suggested by previous experimental data, while the negative feedback and feedforward loops in red are suggested by the cell depletion experiments presented here. See Methods for a detailed description of the corresponding ordinary differential equations. Although our data were derived from mice, we hypothesize that similar cell-cell signaling occurs in humans.

### Parameter estimation for feedback-feedforward model of hematopoiesis

To determine biologically relevant parameters for the feedback-feedforward model in **Fig. 2F**, a grid-search algorithm was employed. The full ODE model is given in Methods and in Supplemental Material (**Sec. S2**). The twelve model parameters (proliferation and death rates, self-renewal and branching fractions, feedback/feedforward gains) were sampled using a random uniform distribution for each parameter. See Methods and Supplemental Material (**Sec. 3, Tables S2, Table S3**) for details and a full parameter list. Once parameter values were chosen, the model was simulated for long times. If a parameter set resulted in steady state values consistent with the range of values previously reported for a dynamic human hematopoiesis model (*Manesso et al., 2013*), that parameter set was accepted. Out of ∼10^6^ possible parameter combinations, a total of 1493 parameter sets were accepted (Supplemental **Fig. S4)**. We further restricted the candidate parameter sets by considering only those with sufficiently large feedforward gains on the MPP division rate (γ_5_>0.01). This reduced the number of eligible parameter sets to 563 and their distributions are shown in Supplemental **Fig. S5**. Each of these parameter sets can be thought of as representing the “normal” condition of a virtual patient by having different individual parameters, e.g., due to genetic, epigenetic or environment factors, that nevertheless result in a “normal” homeostatic hematopoietic system. The different parameter sets thus model a range of variability across individual CML patients. The parameter sets used in **Figs. 3-6** in particular are listed in Supplemental **Table S4**.

**Figure 3:**
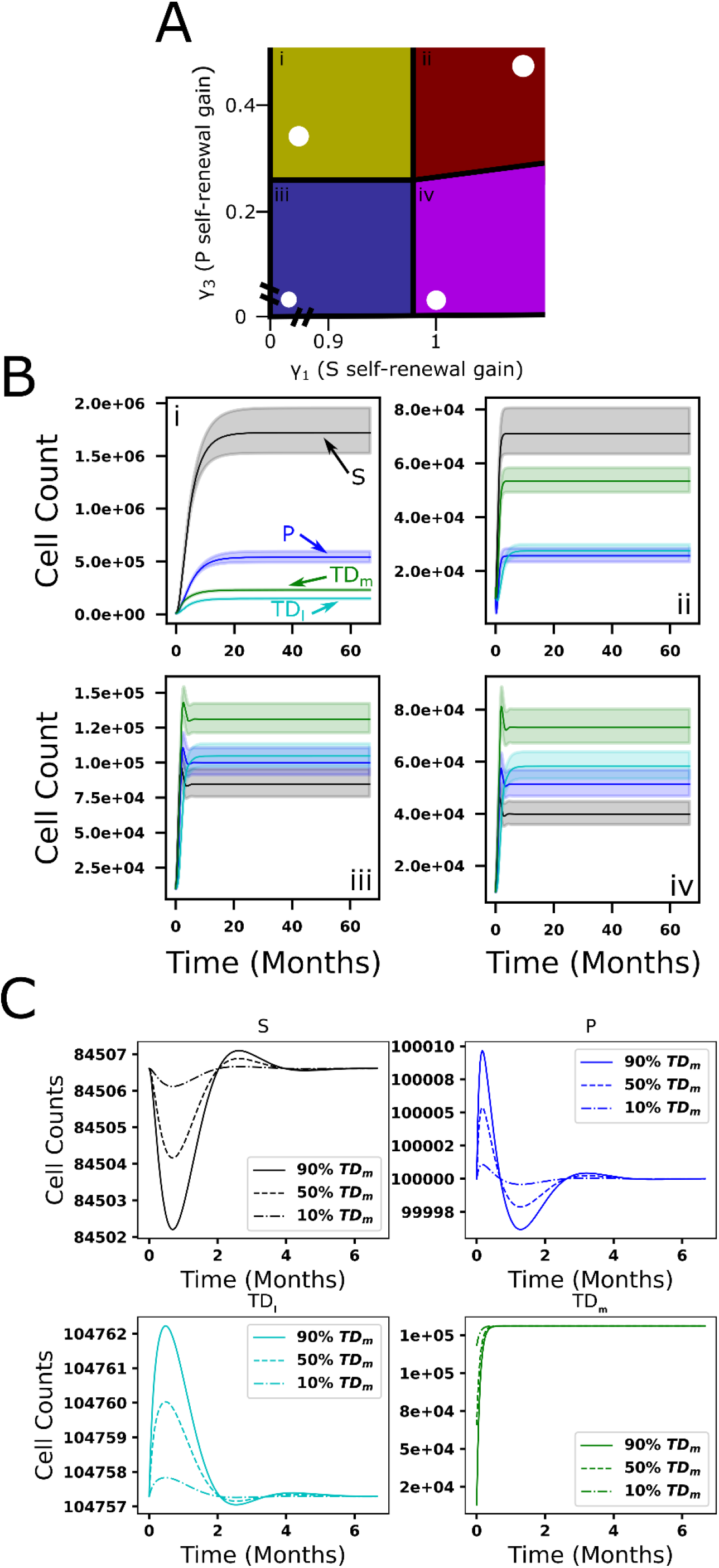
Qualitative behavior of feedforward-feedback model and parameter sensitivity. **A**. The colored regions (i-iv) represent areas of design space in which there are distinct qualitative behaviors as a function of the feedback gains γ_1_ and γ_3_ of the HSC and MPP self-renewal fractions, respectively. White dots denote specific parameter combinations. **B**. The dynamics for each cell compartment within each of the four design space regions (i-iv). Solid lines represent ODE solutions using the specific parameter combinations (black dots in **A**) while the lightly colored regions represent the range of ODE solutions resulting from perturbations in γ_1_ and γ_3_ in a range within 0.9-1.1 times their original values. The blue and black curves correspond to the HSC and MPPs, respectively, the green and turquoise curves correspond to the myeloid and lymphoid cells. **C**. The return to equilibrium following partial depletion of mature myeloid cells (10%, 50%, 90%) using the parameter combination (white dot) in region iii.

**Figure 4:**
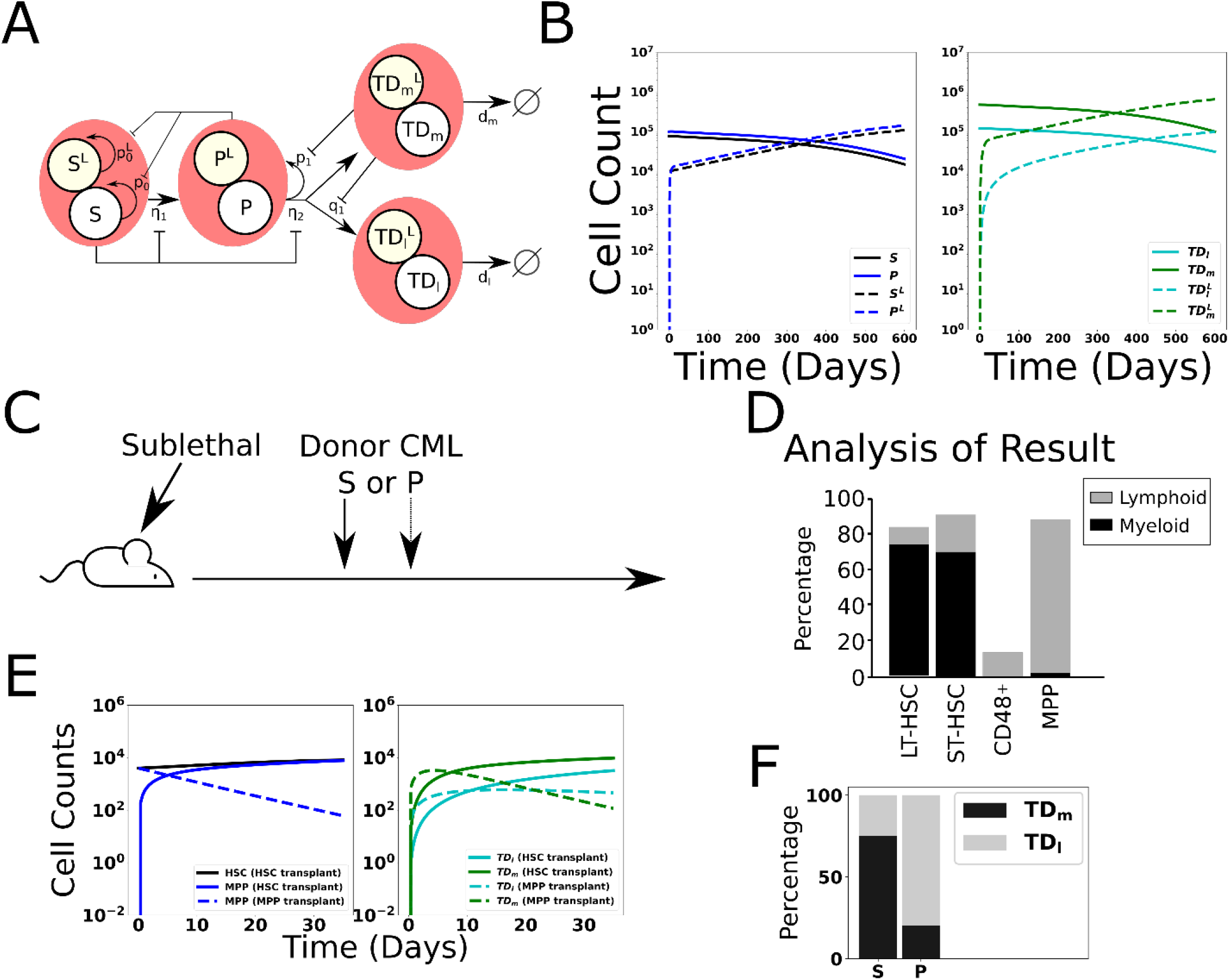
Extension of the model of hematopoiesis to chronic myeloid leukemia. **A**. Schematic of two branched lineages consisting of normal and CML cell compartments. The two lineages share the same feedback architecture. The difference between the two lineages is the leukemic HSC self-renewal is less affected by negative feedback, denoted by p_0_^L^ (see text). **B**. Dynamics of hematopoiesis upon introduction of CML cells. We begin with having normal hematopoiesis at equilibrium. At time 0, 10^4^ leukemic stem cells (HSC^L^, S^L^) cells are introduced to the system and subsequently expand over time at the expense of the normal cells, which decrease. **C-D**. Results of a transplant experiment from Reynaud et al. (*Reynaud et al., 2011*). Schematic (**C**) depicting the experimental pipeline and results (**D**), adapted from (*Reynaud et al., 2011*). When HSC^L^ are transplanted into sublethally irradiated mice, CML-like leukemia is induced and the myeloid cells expand. When leukemic multipotent progenitor cells (MPP^L^, P^L^) are transplanted, they do not stably engraft and transiently produce a larger fraction of differentiated lymphoid cells. **E**. Simulated time evolutions of donor-derived HSC^L^, MPP^L^, and terminally differentiated lymphoid (TD_l_), and myeloid (TD_m_) cells when HSC^L^ (solid) or MPP^L^ (dashed) cells are transplanted. **F**. Bar chart showing model predictions of the percentages of donor-derived myeloid and lymphoid cells after 35 days when HSC^L^ or MPP^L^ are transplanted, which is consistent with the experimental data in **D** (see text).

**Figure 5:**
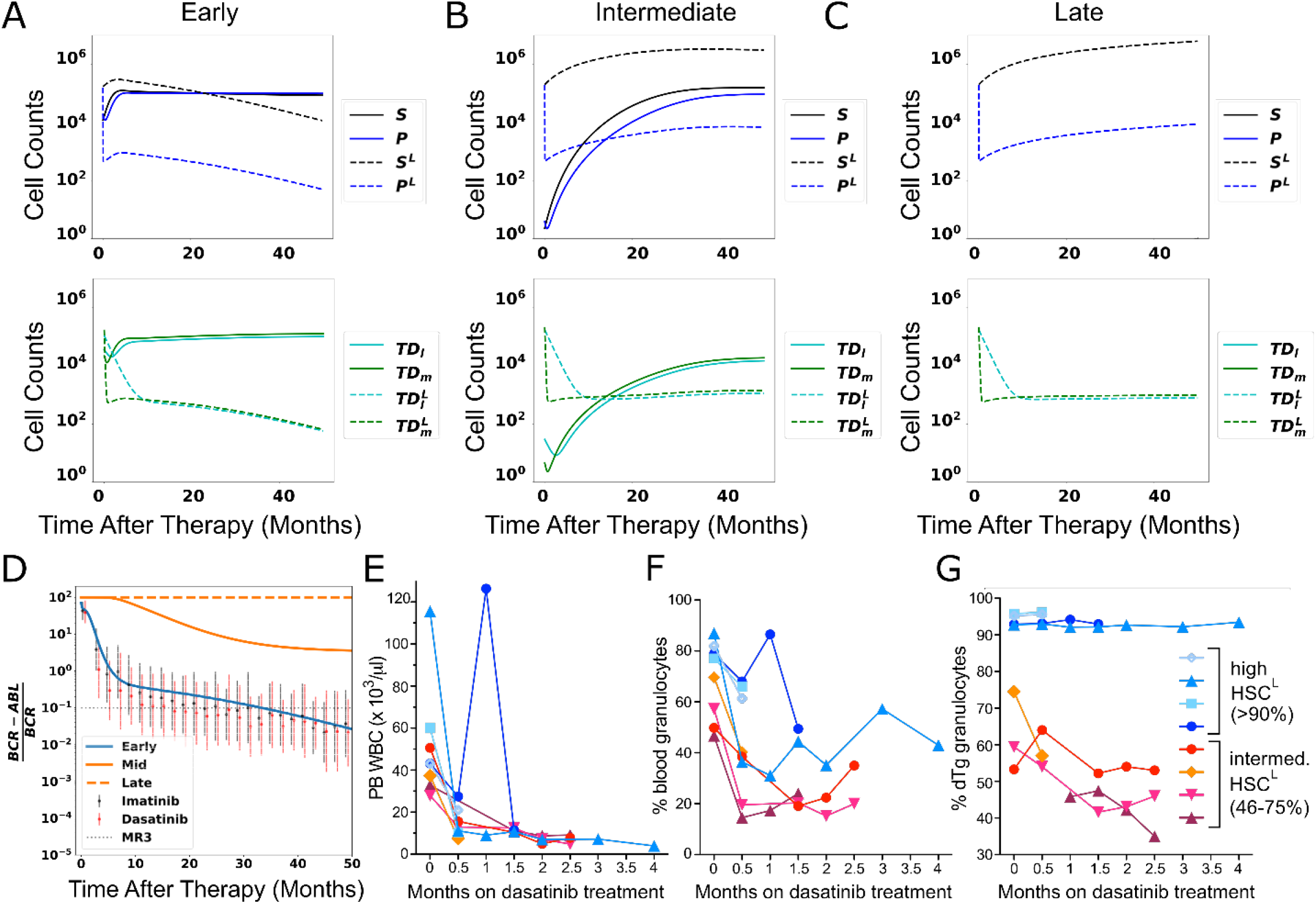
The response of CML to TKI therapy. **A-C**. Simulated cell dynamics of normal and leukemic cells in response to TKI therapy that is started at different time points in CML development (A-early times; B-intermediate times; C-late times). See text. **D**. Simulated molecular response curves corresponding to the application of TKIs for each of simulations in A-C. The simulated molecular response from A (blue) compares well with clinical data (symbols) measuring treatment responses to two different TKIs (imatinib, dasatinib) averaged across a cohort of patients (*Glauche et al., 2018*). The simulated molecular responses from B and C (orange solid and dashed curves) are indicative of primary resistance. **E-G**. Experiments in chimeric mice (see text) that show that the size of the leukemic stem cell clone correlates with decreased response to TKI therapy. Peripheral Blood (PB) leukocyte counts (**E**), percentage of PB granulocytes (**F**), and PB *BCR-ABL1*^+^ (leukemic) granulocyte chimerism (**G**) are shown in cohorts of mice treated with dasatinib. Blue symbols are chimeras bearing >90% *BCR-ABL1*^+^ HSC^L^, red-orange symbols are chimeras bearing 46-75% *BCR-ABL1*^+^ HSC^L^.

**Figure 6:**
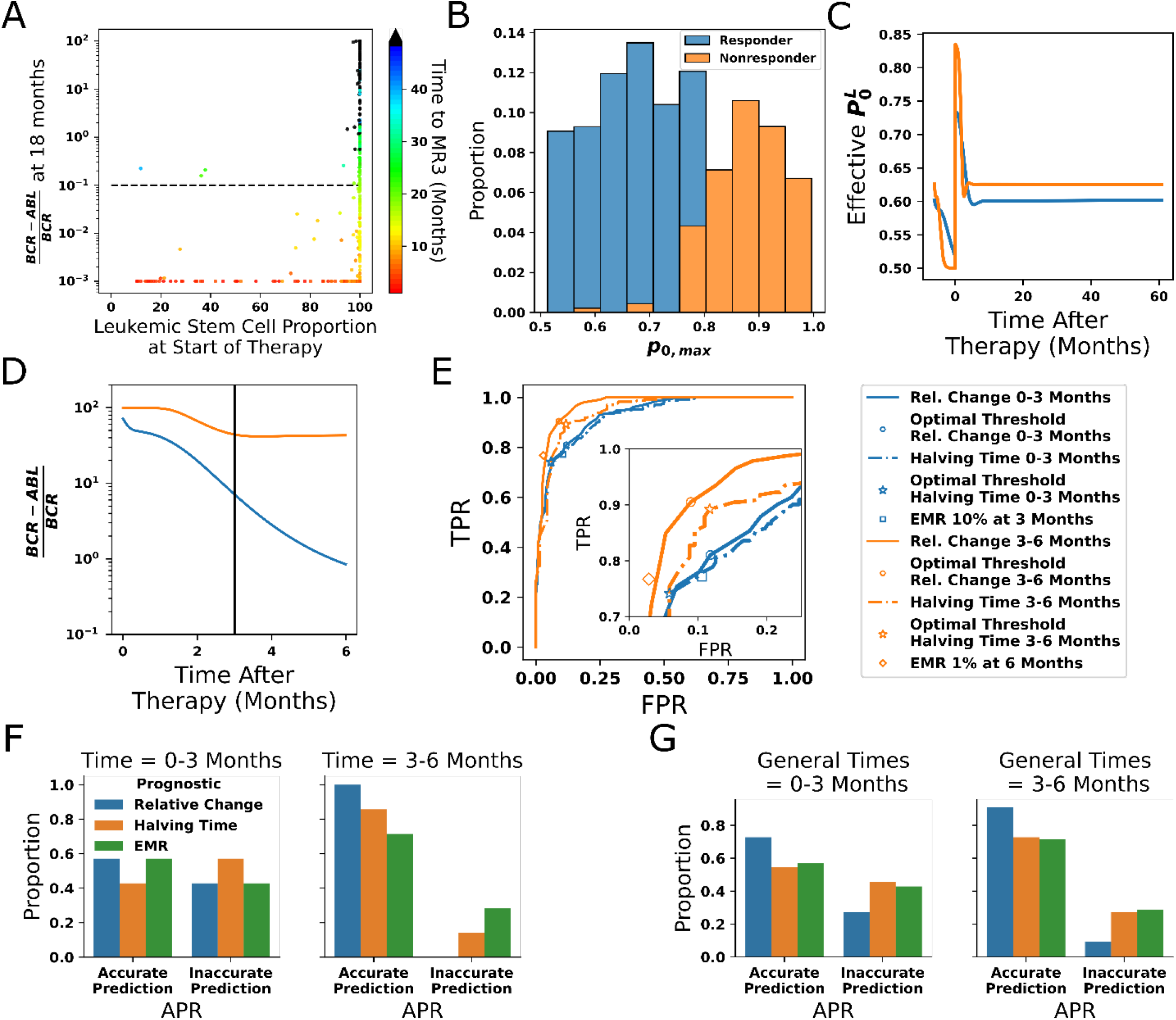
Leukemic stem cell load alone does not predict response to TKI therapy. **A**. Scatter plot of the distribution of simulated *BCR-ABL1* transcripts at 18 months after start of treatment as a function of the HSC^L^ proportion at the start of TKI therapy for each of the 478 parameter sets. The time to reach MR3 (*BCR-ABL1* transcripts less than 0.1%) is indicated by the color. **B**. The maximum stem cell self-renewal fraction distinguishes between parameter sets that achieve MR3 within 50 months (responders, blue) and those that do not (non-responders, orange); the y-axis indicates the proportion of parameter sets for each value of p_0,max_. **C**. Dynamics of the effective leukemic stem cell self-renewal fraction for the parameter set used in Fig. 5A-D (blue) and an arbitrary representative non-responsive parameter set (orange) during CML development and before initiation of therapy (t < 0), and after application of TKI therapy (t > 0). **D**. Early time dynamics (e.g., t = 0 to 3 months; left of the vertical line) of the transcript levels reveal that it is difficult to distinguish responders (blue) from non-responders (orange). At later times (e.g., t=3 to 6 months; right of the vertical line), the two populations are easier to distinguish. **E**. Receiver operating curves (ROC) obtained from the 478 parameter sets using our new prognostic criterion based on the relative changes in transcript levels (solid) and the transcript halving time (dashed) for the first three months (blue) and the second three months (orange) after therapy. The prognostic thresholds (symbols) are identified by optimizing true and false positives rates. EMR at three months (10% transcript levels) and six months (1% transcript levels) are shown by the blue square and orange diamond, respectively. Inset: Expanded view of the “elbow” region of the ROC curves to display differences between the prognostic tests. Accuracy is improved using the 3-6 month time window, and our new prognostic criterion outperforms the EMR and halving time prognostics in this time window. **F**. The accuracy of the prognostic criteria applied to CML patient data treated using the same TKI dosing for the six month period after therapy is started. The results are consistent with the synthetic data in **E. G**. The prognostic criteria applied to patient data in which therapy could be changed but those changes were maintained for six months (see text). Although the data are limited, the results are consistent with those in **E** and **F**, suggesting increased accuracy using the 3-6 month window, and that the prognostic criterion based on relative change may yield more accurate predictions than EMR and halving time in the 3-6 month time frame.

### Sensitivity analyses of hematopoiesis model

DSA can be used to determine qualitative model behaviors and how sensitive the model is to perturbations of key parameters. Here, we focused on the feedback gains γ_1_ and γ_3_ on the HSC and MPP self-renewal probabilities, respectively (see Supplemental Material **Sec. S1.3** for details, and for sensitivity analyses for other parameters, see Supplemental Material **Sec. S4, Fig. S6**). As indicated in **Fig. 3A**, DSA identifies four regions (design space) in the γ_1_ and γ_3_ plane which the dynamics are governed by different S-systems. Using a parameter set in each design space region (indicated by white dots) as a base value, we performed a parameter sweep in which we vary γ_1_ and γ_3_ in a range within 0.9-1.1 times the magnitude of their original values. In **Fig. 3B**, the evolution of each of the cell populations is shown, starting from an initial condition in which there are only a small number of HSC. The different graphs correspond to the parameter sets (Supplemental Material, **Table S4**) in the four regions of the design space although the dynamics are shown for the full ODE solutions. The solid curves denote results from the original (white dot) parameter set and the shading denotes the range of behaviors when the parameters are varied. The black and blue curves correspond to the HSC and MPPs, respectively, while the dark-green and light-green correspond to the terminally-differentiated myeloid and lymphoid cells. While the system tends to equilibrium for all parameter combinations, the approach to equilibrium is different. The dynamics in regions i and ii are monotonic while those in regions iii and iv are not (e.g., the equilibria in regions i and ii are stable nodes, while those in regions iii and iv are stable spirals). Further, the larger the γ_1_, the faster the approach towards equilibrium. The cell numbers and proportions in each design space region are different as well. In regions (i) and (ii), the HSC dominate while in regions (iii) and (iv) the differentiated myeloid cells dominate the population. Further, the number of cells in regions (i) and (iii) is larger than those in regions (ii) and (iv). The equilibrium cell populations in region (iii) correspond more closely to the physiological populations identified by Manesso et al. (*Manesso et al., 2013*).

We next investigated the sensitivity of the model to perturbations about the equilibrium cell population. In **Fig. 3C**, we present results obtained by reducing the number of terminally differentiated myeloid cells from their equilibrium value by 10% (dot-dashed), 50% (dashed) and 90% (solid) and with parameters from design space region iii (Supplemental Material, **Table S4**). By initially depleting the myeloid cells, which is similar to the experiment in **Figs. 2D-E**, the hematopoietic system is shifted away from its steady state. While the presence of the negative feedback loops introduces small magnitude oscillations of the HSC, MPPs and lymphoid cells, the myeloid cell dynamics are monotonic and the system robustly returns to its steady state over times that are consistent with those established in previous experiments (*Reynaud et al., 2011*) for similar perturbation studies.

### Extension of the hematopoiesis model to CML

Following previous modeling studies, we modeled CML by introducing a parallel lineage of mutant leukemic cells (denoted by the superscript L) but with the behavior of that lineage coupled at many points to the behavior of non-mutant cells, and vice-versa. In particular, the model for normal and CML cells share the same lineage structure and feedback architecture with both normal and mutant cell types providing a source of regulating factors. For simplicity, we assumed the only difference between the two lineages is a decrease in the feedback strength for leukemic HSC (**HSC**^**L**^; **S**^**L**^), as indicated by p_0_^L^ in the schematic in **Fig. 4A**. This makes the leukemic cells less responsive to negative feedback and enables leukemic cells to gain a competitive advantage for growth. One candidate mediator of this negative feedback is CCL3, previously shown to inhibit self-renewal and division of normal HSC but HSC^L^ are less sensitive to its inhibitory regulation (*Eaves et al., 1993; Dürig et al., 1999; Baba et al., 2016; Staversky et al., 2018*). An example of CML hematopoiesis is shown in **Fig. 4B**, where it is seen that, after the introduction of a few HSC^L^ at equilibrium of the normal hematopoietic system, the CML cells (dashed curves) repopulate the BM at the expense of normal cells (solid curves). Because of negative feedback, the system will eventually reach a new equilibrium consisting solely of leukemic cells. See Supplemental **Table S5** for the leukemic parameter values, and Supplemental Material **Fig. S7** for parameter sensitivity studies of systems containing both normal and CML cells.

### Validation of the CML model

To test whether our mathematical model recapitulates known features of CML biology, we simulated a transplant experiment in a transgenic mouse model of CML (*Reynaud et al., 2011*). In this experiment, either HSC^L^ or leukemic MPPs (MPP^L^) were implanted into sub-lethally irradiated mice (**Fig. 4C**). Transplantation of HSC^L^ enables engraftment and myeloid cell production that leads to CML. On the other hand, transplanting MPP^L^s does not allow for long-term engraftment but results in a larger fraction of donor-derived lymphoid cells after 35 days (**Fig. 4D**). This study presented evidence that IL-6 produced by differentiated myeloid cells reprograms these MPP^L^ progenitors towards a myeloid fate (*Reynaud et al., 2011*). As described in Methods and Supplemental Material **Sec. S5**), we modeled this experiment by reducing the number of cells in equilibrium to mimic the effects of sublethal radiation. We explored a range of possible reductions of HSC^L^, MPP^L^ and differentiated myeloid and lymphoid cells and tracked the outcomes when 4000 HSC^L^ or MPP^L^ were introduced after the decrements from equilibrium. We then discarded those parameter sets that did not yield results consistent with (*Reynaud et al., 2011*). In particular, 85 parameter sets were discarded leaving a total of 478 parameter sets remaining. Characteristic results are presented in **Figs. 4E** and **4F** when the reductions for HSC^L^, MPP^L^ and terminally differentiated cells were 55%, 35%, and 10%, respectively, from their equilibrium values. See Supplemental Material **Fig. S8** for results using other decrements, the removed parameter set criteria (Supplemental Material **Fig. S9**), and Supplemental Material **Fig. S10** for the final parameter distributions. When HSC^L^ are transplanted (solid curves), the donor-derived MPP^L^ (**Fig. 4E**, left) rapidly increased as did the terminally differentiated myeloid and lymphoid cells (**Fig. 4E**, right). Consistent with the experiments, there is a larger fraction of donor-derived myeloid cells than lymphoid cells after 30 days (**Fig. 4F**). In contrast, when MPP^L^ are introduced (dashed curves), their population decreases (**Fig. 4E**, right) because the MPP^L^ do not stably engraft. Concomitantly, there is burst of donor-derived myeloid and lymphoid cells at early times (**Fig. 4E**, right) as the transplanted MPP^L^ differentiate.

The early time dynamics of the myeloid and lymphoid cells depend on the specific values of the MPP^L^ self-renewal (p_1_) and fate control (q_1_) fractions, whose values in turn depend on the number of myeloid cells through negative feedback regulation. In particular, if 1 − *p*_1_ > 2*q*_1_ then more myeloid than lymphoid cells will be produced at early times, as in **Fig. 4E** (right), whereas more lymphoid cells will be produced if 1 − *p*_1_ < 2*q*_1_. In both cases, because the MPP^L^s do not stably engraft and instead differentiate into lymphoid and myeloid cells, we observe that there is a larger fraction of donor-derived lymphoid cells after 30 days (**Fig. 4F**), consistent with the experiments. This occurs because there is a decreasing flux of differentiating cells since there is no stable engraftment and the lymphoid cells are longer-lived (smaller death rate) than the shorter-lived myeloid cells, which have a larger death rate.

### Leukemic stem cell load influences TKI therapy outcomes

We next explored the effects of TKI therapy on CML in the model. While the overall size of the phenotypic HSC compartment is not increased in CML patients (*Jamieson et al., 2004*), the proportion of HSC^L^ in the BM can vary widely across newly diagnosed CML patients from a few percent to nearly one hundred percent (*Petzer et al., 1996; Diaz-Blanco et al., 2007; Abe et al., 2008; Thielen et al., 2016*). We therefore investigated how the HSC^L^ load in the BM affects therapy outcomes. We used one eligible parameter set (see Supplemental **Tables S4, S5**), which is capable of characterizing the normal state of our simplified model of the hematopoietic system. The initial condition was obtained by simulating the progression of CML, analogous to that shown in **Fig. 4B**, prior to initiating therapy. TKI treatment was initiated at three different times to achieve varying leukemic stem cell load (6, 18, and 36 months) and was simulated by introducing a death rate of HSC^L^ and MPP^L^ proportional to their proliferation rates, with the HSC^L^ proliferation rate lower than that of normal HSC (*Jørgensen et al., 2006*). The TKI treatment parameters were the same across the three cases. See Methods and Supplemental Material for details and **Tables S4, S5** for parameter values. Thus, these cases can be thought of as representing the response of one virtual patient to TKI therapy implemented at different times after disease initiation.

At an early treatment time with lower (<90%) initial HSC^L^ fractions (HSC^L^, **Fig. 5A**), the numbers of MPP^L^ (blue-dashed), leukemic terminally-differentiated lymphoid (light-green-dashed) and myeloid (dark-green-dashed) decrease rapidly at the early stages of treatment and are accompanied by a rapid increase of HSC^L^ due to the loss of negative feedback from the MPP^L^. This loss of negative feedback from the MPP^L^ also results in a rapid increase in the number of normal HSC (black solid curves) that subsequently drives an increase in the normal MPPs (blue solid). The increased number of HSC and HSC^L^ decreases their division rates due to the autocrine negative regulatory loop as well as the division rates of the MPPs and MPP^L^ through feedforward negative regulation. This decreases the flux into the terminally differentiated cell compartments (both normal and leukemic) thereby decreasing their numbers at early times. At later times, both the HSC^L^ and MPP^L^ gradually decrease in response to TKI-induced cell death, which drives an accompanying decrease in the leukemic differentiated cells. A small, transient increase in MPP^L^ is observed before the gradual decline. This is driven primarily by the increase in flux into the MPP^L^ compartment by differentiating HSC^L^ although there is also a small contribution from the feedforward regulation of the MPP^L^ division rate, which lowers the effectiveness of TKI therapy on the MPP^L^. Both of these effects are reduced as the HSC^L^ numbers are decreased by TKI therapy. This in turn increases the effectiveness of TKI therapy in killing leukemic cells at later times and enables the normal cells (solid curves) to rebound toward their pre-leukemic equilibrium values.

At intermediate treatment time with larger (90-99%) HSC^L^ fractions (**Fig. 5B**), the responses of the leukemic and normal cells to TKI treatment at early times are qualitatively similar to those observed in the previous case although the effects are more pronounced. The increase in HSC^L^ is much larger than the previous case because there are fewer normal cells to compete with in the BM. This significantly decreases the HSC/HSC^L^ and MPP/MPP^L^ division rates through the negative feedback/feedforward regulation and correspondingly the rates of TKI-induced cell death. Accordingly, at later times the MPP^L^ population rebounds, driven by the flux of differentiating HSC^L^, and eventually the system reaches a state in which both normal and leukemic cells coexist. The stem cell compartment is dominated by HSC^L^ which are largely quiescent, while the multipotent progenitor and terminally differentiated cell compartments have a higher fraction of normal cells. This is consistent with experimental results from mouse models (*Reynaud et al., 2011*)and our own unpublished data. In this scenario, *BCR-ABL1* transcript levels in the peripheral blood are ∼1-9% but the patient would not respond further to TKI treatment and hence would not reach MR3; this has been observed clinically including one of the patients in our study (see below). The small flux of differentiating normal and leukemic stem and progenitor cells, combined with the negative feedback loops on the self-renewal and branching fractions, supports nearly steady populations of differentiated cells.

When given years to develop and a late time to treatment, the HSC^L^ fraction is nearly 100% (**Fig. 5C**), and there are so few normal stem cells that the leukemic cells easily maintain nearly 100% of each cellular compartment even in the presence of TKI therapy. Aside from a short-lived, transient decrease in MPP^L^ (and differentiated leukemic cell) numbers, the leukemic cells are largely unresponsive to TKI therapy because the feedback/feedforward negative regulation of stem and progenitor cell division rates makes these rates so low that the TKIs are largely ineffective in killing the leukemic stem and progenitor cells. As in the previous case, the negative feedback regulation and the small fluxes of differentiating leukemic stem and progenitor cells enables the system to approach a steady state containing only leukemic cells.

In **Fig. 5D**, we plot the simulated *BCR-ABL1* transcript levels over time for the three scenarios. As described in Methods, the transcripts are modeled using a relative ratio of leukemic and normal terminally differentiated myeloid and lymphoid cell numbers. The solid blue curve corresponds to CML using the treatment time from **Fig. 5A**, which responds to TKI therapy. Just as in the clinical data (symbols), the response to TKI therapy produces a biphasic exponential decrease in *BCR-ABL1* transcripts, which decreases below 10^−1^, representing a so-called major molecular response (MMR or MR3), which represents a major goal of therapy in CML as the risk of relapse and leukemia-related death is virtually non-existent once this milestone is achieved (*Hochhaus et al., 2017*). Consistent with previous interpretations, the rapid initial decrease in *BCR-ABL1* transcripts is due to TKI-induced cell death of MPP^L^ and the increase in normal HSC and MPPs, which induce corresponding changes in the myeloid and lymphoid cells (**Fig. 5A**). The long-term, slower depletion of leukemic cells and the stable normal cell populations result in the second phase of the biphasic response. The simulated results compare well with clinical data from the DAISISON study of imatinib versus dasatinib in patients with newly diagnosed CML (*Cortes et al., 2016*) where the data corresponds to mean *BCR-ABL1* transcripts, with standard deviations, adapted from Glauche et al. (*Glauche et al., 2018*) for patients who received imatinib (blue) of dasatinib (red).

The two other curves in **Fig. 5D** correspond to the treatment times from **Figs. 5B** (solid orange) and **5C** (dashed orange). In these cases, the *BCR-ABL1* transcripts do not decrease below the MR3 threshold, indicating that neither of these virtual patients responds adequately to TKI therapy. There is a partial response in patient from **Fig. 5B** as the transcripts initially decrease due to TKI-mediated death of MPP^L^, but this effect soon saturates because the leukemic stem cells are able drive the re-growth and persistence of leukemic progenitor and differentiated cells. For the virtual patient with parameters from **Fig. 5C**, there is essentially no change in the *BCR-ABL1* transcripts when therapy is applied. These behaviors are consistent with those observed in CML patients with primary resistance to TKI therapy (*Zhang et al., 2009; Yeung et al., 2012; Pietarinen et al., 2016*).

### HSC^L^ load influences the response to TKI therapy in a mouse CML model

The fundamental difference between these three virtual patients is the number of leukemic stem cells at the start of therapy, which occurs because treatment is initiated at different times following the development of CML (early— 6 months after CML initiation ∼93% initial HSC^L^ fraction, intermediate--18 months after CML initiation ∼99% initial HSC^L^ fraction, late—36 months after CML initiation ∼99.99% initial HSC^L^ fraction). Our results suggest that the higher the HSC^L^ fraction at the start of therapy, the less effective the therapy. This follows from the feedback/feedforward regulation where increased numbers of HSC and HSC^L^ decrease their own proliferation rates as well as those of the MPPs and MPP^L^ (see **Figs. S11-13** in Supplemental Material for further explorations of feedback/feedforward regulation of parameters). This reduces the effectiveness of TKI therapy as evidence suggests TKIs preferentially target dividing leukemic cells (*Graham et al., 2002; Corbin et al., 2011*) and suggests a mechanism why some patients are destined to do poorly with TKI therapy. To test this hypothesis, we created BM chimeric mice containing both normal and leukemic (*BCR-ABL1*^+^) HSC and treated them with dasatinib, a second-generation ABL1 TKI (**Figs. 5E-G**). As described in Methods, two cohorts of chimeric mice bearing either a high HSC^L^ burden (94 ± 1.5% of the HSC population) or an intermediate HSC^L^ burden (58 ± 12%) were treated with dasatinib (25 mg/kg daily by oral gavage). Both cohorts showed a hematologic response to TKI therapy, with decreased peripheral blood leukocyte counts (**Fig. 5E**) and a decreased percentage of circulating granulocytes (**Fig. 5F**). By contrast and consistent with the predictions of the quantitative model, while mice bearing smaller populations of HSC^L^ showed a decrease in the percentage of circulating *BCR-ABL1*^+^ granulocytes in response to TKI therapy, mice with the highest HSC^L^ burden showed virtually no decrease in circulating leukemic cells (**Fig. 5H**). Because the level of circulating granulocytes reflects the proportion of BM HSC (*Wright et al., 2001*) (and data not shown), these results demonstrate that TKI therapy is unable decrement the HSC^L^ compartment in mice with predominantly *BCR-ABL1*^+^ HSC at the start of treatment.

### HSC self-renewal as an additional determinant of TKI response

While analyses of clinical data also show that patients with lower leukemic stem cell burden are more likely to respond to TKI treatment (*Thielen et al., 2016*), some patients with high percentages of leukemic stem cells at the start of treatment are nonetheless still capable of responding to TKI therapy (e.g., see Fig. 3 in Thielen et al. (*Thielen et al., 2016*)). This suggests that leukemic stem cell burden alone does not predict the molecular response to TKIs. To investigate this further, we tested the response to TKI treatment for each of our 478 parameter sets. The results shown in **Fig. 6A** bear a striking resemblance to the clinical data of Thielen et al. (*Thielen et al., 2016*). The leukemic stem cell fraction does influence TKI response, but treatment outcomes are seen to vary among virtual patients within the same initial leukemic stem cell load. We then asked what characteristics (e.g., parameter sets) distinguish whether a virtual patient achieves a MR3 response within 50 months. Surprisingly, only two parameters were found to clearly distinguish responders from non-responders: the maximal self-renewal fraction p_0,max_ for normal stem cells, shown in **Fig. 6B**, and the negative feedback gain on normal stem cell self-renewal (γ_1_) (see Supplemental Material **Figs. S14** for the distribution of γ_1_ and all the parameter distributions). Larger values of p_0,max_ and γ_1_ are correlated with a decreased response to TKI therapy after leukemia develops. Although these parameters are associated with normal HSC, the self-renewal fraction p_0_^L^ of HSC^L^ also depends on these parameters (see Methods and Supplemental Material **Secs. S2, S3** and **Tables S2, S3**). To compare between response and non-response we take the parameter set from **Fig. 5A-D** as a representative patient for response and select an arbitrary non-responsive parameter set (Supplemental Material, **Table S6**) to be a representative patient for non-response. In **Fig. 6C**, we show that the effective p_0_^L^ (the fraction of HSC^L^ self-renewal after feedback) for non-responders (orange) is larger after TKI therapy is applied than for responders (blue). In particular, as TKI treatment kills the leukemic progenitors, this increases the effective self-renewal fraction for both normal and leukemic stem cells because of the release of negative feedback. When the maximum self-renewal p_0,max_ is larger, the leukemic stem cells experience a higher spike in self-renewal, resulting in their dominance over normal stem cells that then leads to a decreased response to TKIs.

Clinical data provides support for this mechanism of resistance. Patients with clonal hematopoiesis, in which there is a dominant clone driving hematopoiesis, exhibit predominantly normal hematopoiesis but frequently have mutations in the genes, such as *TET2, DNMT3A*, and *ASXL1* that are known to increase stem cell self-renewal (*Steensma, 2018*). Clinical data shows that mutations in *TET2* and *ASXL1*, which may exist prior to development of CML, are also associated with a poor response to TKI therapy (*Kim et al., 2017; Marum et al., 2017*). Taken together, these data suggest that patients who exhibit normal hematopoiesis but with higher stem cell self-renewal fare worse when their CML is treated using TKIs than patients with lower stem cell self-renewal.

### Predicting long-term response to TKI treatment

Several measures of the response of CML patients to TKI therapy have been developed, based on *BCR-ABL1* transcript levels in peripheral blood. Here we test a new, model-driven criterion for predicting patient response and compare the results with several other criteria currently used in the clinic. A major focus has been on the predictive value of the decline in transcripts over the first 3 months of treatment, principally the so-called “early molecular response” or EMR (defined as *BCR-ABL1* transcripts <10% at 3 months and <1% at 6 months), where patients with >10% transcripts had significantly lower probability of achieving cytogenetic remission and decreased overall survival (*Hanfstein et al., 2012; Marin et al., 2012*). Subsequently there was an effort to improve the predictive power by focusing on the velocity of reduction in transcripts (*Branford et al., 2014; Hanfstein et al., 2014; Pennisi et al., 2019*). Because the best predictor of patient response to TKIs, the self-renewal fraction of normal stem cells, is very difficult to measure clinically, we searched for an alternative criterion that could accurately predict patient response and could still be measured using data collected in standard practice. Therefore, we focused on alternative time frames and calculation methods for assessing *BCR-ABL1* transcript levels (**Fig. 6D**). It is important to be able to predict the long-term TKI response early after starting treatment in order to enable changes in therapy. However, since both responders (blue) and non-responders (orange) may show significant decreases in the transcript levels in the first months of treatment, it was difficult to distinguish between the two at relatively early time points. By contrast, responses in the 3-6 month time frame make it easier to identify the different behaviors of responders and non-responders (**Fig. 6D**).

By calculating the relative changes of the transcript levels from 3 to 6 months, we developed a prognostic formula: *PF*(3, 6) = *BCRABL*1(6) − *BCRABL*1(3)/*BCRABL*1(6). We found that optimizing for sensitivity (TPR, the True Positive Rate) and specificity (1-FPR, with FPR being the False Positive Rate) resulted in a prognostic threshold of *PF* ≈ −3.2, with sensitivity of ≈ 0.91 and specificity of ≈ 0.91 (**Fig. 6E, orange curves**) compared to the optimal sensitivity and specificity of the velocity-based prognostic (≈ 0.89 and ≈ 0.88, respectively) and ≈ 0.77 and ≈ 0.98 for EMR 1% with our parameter sets. This demonstrates that this prognostic tool had higher sensitivity and specificity than previously developed predictive criteria in separating responders (< −3.2) from non-responders (> −3.2), where response is defined as achievement of MR3 within a clinically relevant timeframe of 18 months. We also tested the various prognostics at the 0-3 month interval as is the current clinical practice, but that resulted in lower predictive power (**Fig. 6E**, blue curves). These results highlight the importance of including the 3 to 6 month TKI response in predicting the long-term outcome of treatment, instead of considering only the first three months. See Supplemental Material (**Fig. S15**) for further discussion and comparison of additional prognostic criteria.

We then applied our prognostic criterion to anonymized CML patient data (see Methods) to determine clinical significance and utility. The prognostic tests shown in **Fig. 6E** were calculated for both the first three months and the subsequent three to six month period after the start of therapy, for the patients who were treated with the same TKI and dosage for the full six months. All the prognostic tests achieved a more accurate prediction of patient outcome using the three to six month data compared to the same test applied to the first three months (**Fig. 6F**). To expand clinical utility, the prognostics were calculated for cases where TKI therapy was changed (due either to toxicity or an inadequate response) but then maintained for a subsequent 6 month period, which were added to the data from **Fig. 6F**. The aggregated data (**Fig. 6G**) re-affirms the improved accuracy in prediction using the three to six month transcript data compared to that from zero to three months. Over the first three months, all the prognostic criteria performed similarly. Although the number of patients was limited, the results suggest that our prognostic criteria may perform better than the EMR and velocity-based prognostics that are in current clinical use. For comparisons between the prognostic criteria and time frames see Supplemental Material **Sec. S8, Figs. S15-18**.

### Improving response to therapy: Combining TKIs with differentiation promoters

Our model suggests that combination therapy to modulate the stem cell self-renewal rate in addition to directly targeting the leukemic HSC and MPPs with TKI therapy might counteract TKI treatment resistance mediated by high stem cell self-renewal. Such pro-differentiation therapy could be accomplished through either direct stimulation of differentiation or through suppression of self-renewal. In our modeling experiments, we explored differentiation therapy through the suppression of self-renewal (see Methods for details). To begin the exploration of the combined TKI-differentiation therapy, we performed this combination therapy on each of our 478 parameter sets, which represents a population of CML patients with person-to-person variability. We then recorded which parameter sets achieved MR3 within 50 months for each strength of the differentiation therapy (Δ), where Δ is a dimensionless constant greater than 0 that constantly suppresses stem cell self-renewal (both normal and CML) in the setting of combination therapy (see Methods and Supplemental Material **Eqns. S40**-**S41**). Using these data, **Fig. 7A** depicts the proportion of parameter sets achieving MR3 given a strength of differentiation therapy of Δ. As differentiation therapy strength increases from zero, the proportion of parameter sets that achieve MR3 increases before leveling off between Δ=0.2-0.3, with maximum efficacy occurring at a strength of differentiation Δ of about 0.29. The efficacy of combination therapy then begins to decline rapidly, and with too great a strength of differentiation treatment, the combination therapy becomes inferior to TKI therapy alone.

**Figure 7:**
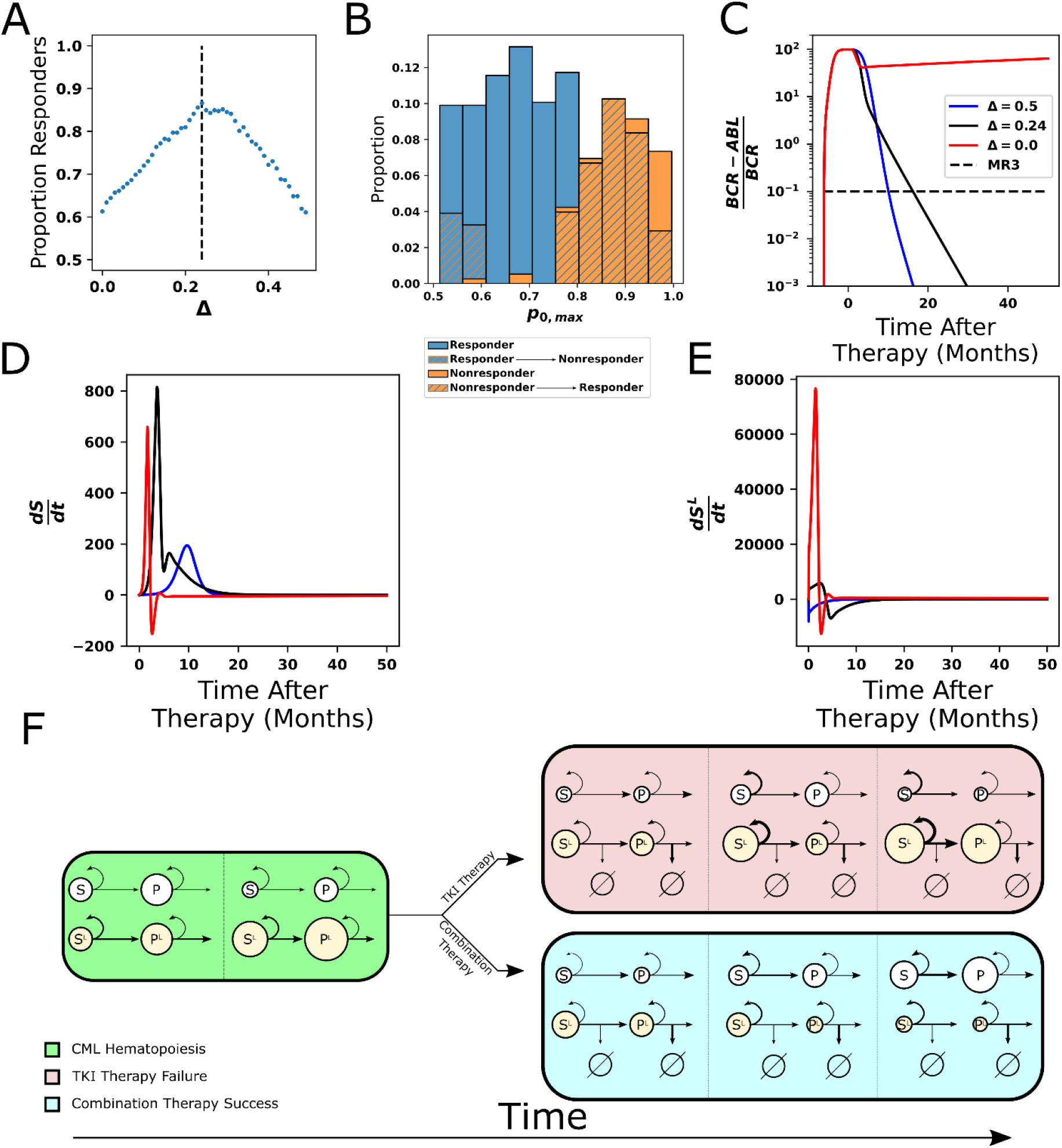
Combining TKI therapy with differentiation promoters enhances response to treatment. **A**. The proportion of the 478 parameter sets that achieve MR3 under combined TKI and differentiation therapy depends non-monotonically on the strength Δ of the differentiation therapy, with the peak response (86.8%) occurring at Δ=0.29. **B**. The maximum stem cell self-renewal fraction from Fig. 6B with hatching indicating the effects of the combination of TKI and differentiation therapy with Δ=0.29. Blue hatching indicates non-responders (who did not achieve MR3) that become responders (achieve MR3) while orange hatching indicates responders that become non-responders upon combined treatment. Differentiation promotors allow non-responders to TKI therapy with large self-renewal fractions to reach MR3. The opposite outcome, loss of MR3 in a TKI responder, primarily occurs only at the smallest self-renewal fractions. **C**. Time evolution of *BCR-ABL1* transcripts during combination therapy, with Δ=0.29 (black) and Δ=0.5 (blue), using the parameter set from Figure 5B that does not achieve MR3 using TKI monotherapy (red). **D-E**. The time derivatives of the number of normal (**D**) and leukemic (**E**) stem cells during combination therapy. The differentiation promoter attenuates the rapid increases in the rates of change at early times after therapy starts in both normal and leukemic cells, but the attenuation is much larger in the leukemic cells. This results in the growth of normal cells, while leukemic cells experience restricted growth or outright depletion depending upon the differentiation therapy strength. **F**. Simplified diagram representing the key interactions between the cells and the impact on outcomes of TKI and combination therapy. Green: CML hematopoiesis depicting the loss of normal stem cells and progenitors and the increase of leukemic stem cells and progenitors. Red Fill: TKI treatment failure. The TKI-induced death of leukemic progenitors relieves negative feedback and increases stem cell self-renewal resulting in increases in both normal and leukemic stem cells, and eventually their progeny (panel 2). The increases are larger for the leukemic cells because their self-renewal fraction is bigger. Increases in the leukemic progenitor compartment (panel 3) drive down the self-renewal fraction of normal stem cells proportionally more than for the leukemic stem cells. The increases in HSC^L^ also drive down proliferation rates, which makes the leukemic cells less responsive to TKI treatment. Altogether, this makes the leukemic cells more fit than the normal cells and results in therapy failure. Blue Fill: Treatment by combined TKI and pro-differentiation therapy reduces stem cell self-renewal relative to TKI monotherapies, equalizes the normal and leukemic self-renewal fractions, which limits leukemic stem cell growth and limits decreases in proliferation rates, making the HSC^L^ and MPP^L^ more susceptible to TKI-induced death (panel 2). This allows repopulation of the bone marrow by normal stem cells and progenitors to occur (panel 3).

To investigate how combination therapy effectively targets resistance, and the mechanism of the decreased efficacy of combination therapy in achieving MR3 when Δ is large, we returned to examining parameter distributions. **Fig. 7B** depicts the same distribution of p_0,max_ as in **Fig. 6B**, but overlaid with a second histogram (hatched regions) to denote the effect of the differentiation therapy at the point of maximum efficacy (see Supplemental Material **Fig. S19** for all the parameter distributions). The two types of hatching reveal important factors that determine under which conditions combination therapy improves or impairs response. The orange hatching represents transition from response to nonresponse by combination therapy; this occurs in individuals with the lowest p_0,max_. In these cases where stem cell self-renewal is already close to the ideal effective self-renewal fraction of 0.5, differentiation therapy pushes too many normal cells into differentiation, causing the normal cell populations to deplete themselves and decreasing the efficacy as Δ increases beyond 0.29. In contrast, the blue hatching shows the desired scenario of non-responding individuals with high p_0,max_ becoming responders and achieving MR3 within 50 months due to the combination therapy.

To understand further the mechanisms underlying the efficacy of combination therapy, we explored treatment dynamics (changes in *BCR-ABL1* transcript levels) and the rates of change of both normal and leukemic stem cell populations for the nonresponsive individual from **Fig. 6D**. By applying two different strengths of differentiation therapy (Δ = 0.29 and 0.5) in combination with TKI therapy, for this individual both strengths are able to achieve MR3 at ∼18 months (**Fig. 7C**) in contrast to TKI monotherapy, which resulted in a failure to reach MR3 (**Fig. 6B**, orange). **Figs. 7D, E** show how the rates of change in the size of the normal and leukemic stem cell compartments vary with respect to time for the three different Δ values. For TKI therapy alone (Δ = 0), rates of growth of both the normal and leukemic stem cell populations show an increase as a result of the loss of negative feedback due to TKI-induced killing of MPP^L^, but the leukemic stem cells experience a much greater numerical increase and outcompete normal cells resulting in a system that exhibits resistance to the TKI therapy. Under conditions of maximum efficacy (Δ =0.29), the normal stem cell population rate of change still increases rapidly but to a maximum level below that for TKI monotherapy before decreasing more rapidly to zero as the system re-equilibrates. In contrast, under the same conditions the rate of change of the leukemic stem cell population is greatly reduced and becomes negative after normal stem cells begin to outcompete the leukemic stem cells. Under conditions of stronger differentiation therapy (Δ =0.5), although the accumulation rate of the normal stem cells is substantially reduced, the growth rate of the HSC^L^ is immediately negative. This enables the normal HSC to easily outcompete the leukemic cells and restore the system to the normal state. Effectively, for large values of HSC self-renewal and corresponding feedback gains, the differentiation promoter acts to bring the self-renewal fraction of the normal HSC closer to that of the HSC^L^, which then enables the TKI therapy to disadvantage the leukemic cells, and allow for repopulation and dominance by normal cells.

## Discussion

In this work, we developed a nonlinear mathematical model of normal and CML hematopoiesis that incorporated feedback control, lineage branching and signaling between normal and CML cells. Using ODEs, we modeled the dynamics of the stem, multipotent progenitor and terminally differentiated cell populations. To filter through the combinatorial explosion of models that occurs when cell-cell signaling interactions are taken into account, we focused first on normal hematopoiesis. We used design space analysis (*Savageau et al., 2009; Fasani and Savageau, 2010; Lomnitz and Savageau, 2013; Lomnitz and Savageau, 2016*), an approach that enables models to be distinguished based on their range of qualitatively distinct behaviors without relying on knowledge of specific values of the parameters, to perform an automated search for regions of stability in thousands of proposed models and efficiently eliminate unphysiological, unstable models. When combined with previous observations and new *in vivo* data to further constrain cell-cell interactions, we arrived at a new feedback-feedforward model.

We used a grid search algorithm to determine a set of approximately 500 biologically relevant parameter sets for our new model. In particular, using each parameter set in the model yields steady states that are consistent with normal ranges of hematopoietic cells. These parameter sets model a population of individuals with normal cell counts but person-to-person variability of parameters due, for example, to genetic, epigenetic and/or environmental differences.

We then extended the model to incorporate CML hematopoiesis by introducing a mutant lineage with the same structure as the normal system. We incorporated one of the central features of CML pathophysiology, that the leukemic stem cell clone, hypothesized to arise from a single HSC that acquires a Ph chromosome, has a competitive advantage over normal HSC and with time comes to dominate the stem cell compartment (*Dingli et al., 2010; Thielen et al., 2016; Holyoake and Vetrie, 2017; Majeti et al., 2022*). This competitive advantage could be a consequence of positive feedback (autocrine or paracrine) on the HSC^L^ population, or negative feedback with different strengths for normal and leukemic stem cells. Candidate mediators of such positive and negative feedback include interleukin-3 (*Jiang et al., 1999*) and CCL3 (*Baba et al., 2016*), respectively. Our current model incorporated differential negative feedback of MPPs on HSC (**Fig. 4A**) with the HSC^L^ being less sensitive to the negative feedback than are the normal HSC, which is consistent with CCL3 (*Eaves et al., 1993; Baba et al., 2013*). This one difference provided leukemic cells with a competitive advantage for growth, and in the absence of treatment, the leukemic cells will take over the BM at the expense of normal cells (**Fig. 4B**).

When combined with TKI therapy, the feedback/feedforward model exhibited variable responses to TKI treatment, consistent with those observed in CML patients. That is, although our 500 parameter sets were consistent with normal hematopoietic cell counts, the responses to TKI treatment were highly variable, with some sets responding to treatment while others did not. The model predicted that a contributor to primary TKI resistance is the overall proportion of HSC that are leukemic, consistent with experimental data in mice (**Fig. 5G**) as well as patient data (*Thielen et al., 2016*). However, leukemic stem cell burden alone does not predict the molecular response to TKIs, as observed both clinically (*Thielen et al., 2016*) and in our data (**Fig. 6A**), since some patients with high HSC^L^ fractions in their bone marrow nonetheless still respond to TKIs.

The model suggested that a key predictor of reduced response to TKI treatment is an increased tendency of normal hematopoietic stem cells to self-renew, which in turn influences self-renewal of the leukemic stem cells. This is consistent with clinical data that suggest that CML patients with pre-existing mutations in genes such as *TET2* and *ASXL1*, which are known to increase stem cell self-renewal (*Steensma, 2018*), tend to have inferior outcomes under TKI therapy (*Kim et al., 2017; Marum et al., 2017*). This is illustrated in **Fig. 7F** (red panel), where the high initial HSC^L^ population and the subsequent decline of progenitor cells reveals the effect that high stem cell self-renewal has on driving TKI resistance. This self-renewal-driven resistance challenges the prevailing paradigm that TKI resistance is proliferation-driven and a consequence of HSC^L^ quiescence (*Graham et al., 2002; Corbin et al., 2011*).

Because stem cell self-renewal is hard to quantify experimentally, we developed a clinical prognostic criterion to predict TKI response based on the relative changes in the *BCR-ABL1* transcripts over a three month period. Using the synthetic data from our 500 parameter sets, we found that using changes in transcripts from three to six months was very effective in predicting the long-term outcome of treatment (e.g., reaching MR3 within 18 months). In contrast, using transcript data from zero to three months resulted in less accurate predictions. This observation also holds for prognostic criteria based on EMR and transcript halving time, which are currently used in the clinic. We then tested the prognostic criteria on data obtained from small number of anonymized CML patients and found the same conclusions hold. Our results suggest that the relative change prognostic criterion more accurately predicts patient response than EMR and the halving time, although more data are needed to confirm this.

Two strategies can be postulated to overcome TKI resistance. One approach could be to decrease stem cell self-renewal, either by inhibiting pathways implicated in self-renewal (such as those downstream of the epigenetic regulator *TET2*) or by promoting differentiation directly (for example, using retinoids (*Drumea et al., 2008*)). By applying combined TKI and pro-differentiation therapy, the self-renewal fractions of the normal and leukemic stem cells can be decreased and brought closer together, which ultimately disadvantages the leukemic cells because of TKI-induced cell death (**Fig. 7F**, blue panel). An alternative would be to increase stem cell proliferation via pro-proliferation stimuli such as IFN-alpha (*Essers et al., 2009*)to increase efficacy of TKIs in killing HSC^L^.

It is apparent that the feedback/feedforward interactions incorporated in our model, which are necessarily somewhat restricted, may be further constrained by spatial characteristics of the bone marrow microenvironment. Nonetheless, our model still displays consistent and biologically relevant behaviors and although further refinement of the model behaviors is possible, based upon our findings the key behaviors (feedback mechanisms and importance of stem cell self-renewal) would be expected to remain much the same. To explore experimentally observed phenomena not captured by our current model such as treatment free remission, where a low level of HSC^L^ persists in the absence of TKI pressure without myeloid cell expansion, improvement of the model is necessary. For example, it may be necessary to incorporate features of the bone marrow microenvironment such as stem cell-niche interactions (*MacLean et al., 2014; Lai et al., 2022*) and interactions with immune cells (*Hähnel et al., 2020*). The inclusion of a quiescent stem cell state and additional cellular compartments (such as committed progenitors) coupled with appropriately constrained cell-cell signaling would also make the model more physiologically accurate.

In summary, the feedback/feedforward model we have presented here, while a simplified version of normal and CML hematopoiesis, makes novel and testable predictions regarding the origins of non-genetic primary resistance, which patients will respond to TKI treatment and suggests a combination therapy that can overcome primary resistance. Although preliminary evidence was presented to support model predictions, future work should focus on designing targeted experiments and collecting patient outcomes to generate data to more thoroughly test the model.

## Methods

### Mathematical model of hematopoiesis

The classical depiction of hematopoiesis is a hierarchy of cell types starting with the hematopoietic stem cell at the top, followed by progenitors and ultimately ending with mature cells located in the peripheral blood. Therefore, we model hematopoiesis using a lineage ordinary differential equation model (*Roeder et al., 2006; Komarova and Wodarz, 2007; Horn et al., 2008; Foo et al., 2009; Lander et al., 2009; Marciniak-Czochra et al., 2009; Manesso et al., 2013; Buzi et al., 2015; Hähnel et al., 2020; Pedersen et al., 2021*) to describe cellular growth dynamics. The modeling allows us to follow the similar hierarchical structure by creating an order of differentiation. Our branched lineage model of hematopoiesis is simplified and only models hematopoietic stem cells (HSC), progenitor cells (MPPs), and two types of terminally differentiated cells (Myeloid and Lymphoid cells). The model can easily include additional cell types, such as committed progenitor cell types, which will provide an additional level of detail. The model consists of two dividing cell types consisting of S (HSC) and P (MPP) cells with a division rate associated to the cells (η_1_ and η_2_ respectively). The S cells have the ability to self-renew with fractions (p_0_) or differentiate (1-p_0_). The P cells have the ability to self-renew with fraction (p_1_) or differentiate into either TD_l_ (lymphoid) or TD_m_ (myeloid) cells (q_1_ or 1-p_1_-q_1_ respectively). Both S and P cells do not die within the model. The terminal cells form the majority of the hematopoietic system and consist of TD_l_ and TD_m_ cells. TD_m_ and TD_l_ cells are post-mitotic and die at rates d_m_ and d_l_, respectively. The equations below (Eq. 1-4) describe the dynamics of the system:

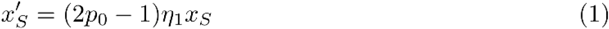

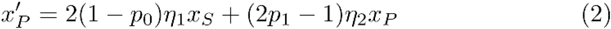

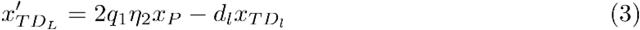

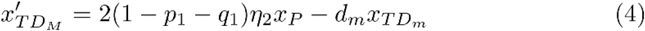

Further expanded forms of the equations are shown in Supplemental **Eqns. 26-29** with the addition of feedback regulation for each of the parameters.

### Design space analysis

We use an automated method developed by Savageau and collaborators (*Savageau et al., 2009; Fasani and Savageau, 2010; Lomnitz and Savageau, 2013; Lomnitz and Savageau, 2016*)that separates models by distinct qualitative behaviors at steady state. The strategy is to deconstruct the model of interest at steady state to focus on cases where one production term and one loss term dominate, which gives a dominant subsystem (S-System). This implies that particular inequalities hold in order to ensure the production and loss terms chosen are larger than the others. The inequalities are evaluated at the S-System’s steady state to assess self-consistency. If the inequalities are satisfied, the system is self-consistent and the regions where equality holds form boundaries that pertain to a particular qualitative behavior associated with the system. The interior region (where strict inequality holds) is termed a domain in design space. If all the S-systems associated with a model do not have any equilibria that are self-consistent or equilibria that are stable, then the model is rejected. The benefits of this method are that it does not require prior knowledge about parameter values, and it can enumerate the different types of qualitative dynamics a certain system may have. By eliminating subsets of parameters for which the equilibrium is unstable, this approach will automatically select models that are robust to parameter variation due to stability. When we applied this method to the ODE system in **Fig. 2F** (supplemental equations (26)-(29)) we found that only the four model classes shown in **Fig. 1B** were accepted. See Supplemental Material **Sec. S1** for further details.

### Parameter estimation

To approximate biologically relevant parameters for the model a grid-search algorithm was employed. Parameters were sampled using a random uniform distribution for each parameter (see Supplemental Material **Sec. S3.1**). Once parameter values were chosen the model was simulated to steady state. If a parameter set resulted in steady state values consistent with the order of magnitude in (*Manesso et al., 2013*), the parameter set was accepted otherwise it was rejected. Specifically, these inequalities had to be satisfied 10^4^<HSC<MPPs with MPPs fixed at 10^5^ and MPPs<TD_l_<TD_m_. For 10^6^ iterations a sample of 1493 parameter sets were accepted. The distribution for these parameter sets is shown in Supplemental **Fig. S4**. To further explore the effect of the feedforward interaction these parameter sets were reduced to the 593 sets with γ_5_>0.01. The distribution for these sets is shown in Supplemental **Fig. S5**. The parameter sets used in **Figs. 3-6** are in Supplemental **Table S4**.

### Modeling CML development

To model CML development in the presence of normal hematopoietic cells we introduce a new leukemic cell type for each compartment. Each compartment is then composed of both normal and leukemic sub-compartments, which exhibit feedback together as a single compartment. We assume the only difference between the two cell lineages is the feedback strength for leukemic HSC self-renewal. This small difference gives the leukemic lineage a competitive advantage for growth, consistent with the ability for leukemic HSC having the ability to initiate CML (*Reynaud et al., 2011; Holyoake and Vetrie, 2017*) and the differential response of the normal and leukemic cells to CCL3 (*Eaves et al., 1993; Baba et al., 2013*), which negatively regulates stem cell self-renewal. The full equations used in the model are shown in Supplemental **Eqns. 30-37**.

### Modeling transplant experiments

The model was tested by simulating the transplant experiments (**Figs. 4C-D**) of (*Reynaud et al., 2011*) where HSC^L^ or MPP^L^ were implanted into sub-lethally irradiated mice and terminal cell counts were measured after 35 days. We used two parallel lineages of leukemic cells with identical parameters to mirror the two leukemic cell populations of the experiment. To mimic the effects of sublethal radiation, we reduced the cell populations from their equilibrium values by variable amounts. The HSC^L^ depletions varied between 50-70% and the MPP^L^ depletions varied between 30-50% while both terminal cells were depleted by 10%. After depletion, an additional 4000 cells of either stem or progenitor types were transplanted in accordance with the experiment. We then discarded the 85 parameter sets that were not consistent with the results from (*Reynaud et al., 2011*), leaving 478 eligible parameter sets. The results shown in **Fig. 4** used depletions of 55% for HSC^L^, 35% for MPP^L^s. See Supplemental Material **Sec. S5** and **Figs. S11-13** for results using other decrements, the discarded parameter sets, and the final parameter distributions.

### Modeling TKI therapy

To account for the treatment by TKIs, additional proliferation-dependent death terms are added to the equations for leukemic stem cells and leukemic progenitor cells shown in Supplemental **Eqns. 38-39** (parameter values are given in Supplemental **Table S5.)**. These represent the ability of TKIs to induce cell death in the leukemic cells. Both cell types have unique death rates, to reflect TKIs having different efficacy in killing stem cells and progenitors. The death rates were selected using a single parameter set to ensure a reasonable biphasic curve for *BCR-ABL1* transcript levels compared to patient transcript levels from (*Glauche et al., 2018*). The same death rates were then used across every parameter set to ensure consistency. In addition to these changes upon initiation of TKI therapy the leukemic stem cell division rate is reduced. This reduction models the ability of TKIs to drive leukemic stem cells to quiescence (*Jørgensen et al., 2006*).

To approximate the *BCR-ABL1* transcript levels we used a method based upon (*Michor et al., 2005*). We use the cell counts of both normal and leukemic terminal cells for both myeloid and lymphoid lineages. The terminal cells are used as in our model they are the closest to peripheral blood in which transcript levels are measured clinically. This results in the following measure for *BCR-ABL1* transcript levels: 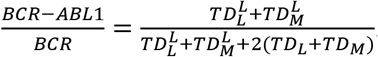.

### Modeling Combined TKI and Differentiation Therapy

Combination therapy consists of simultaneously employing TKI therapy, described in Methods and in Supplemental Material **Sec. S2** and **S3**, and the addition of a new differentiation therapy. To model differentiation therapy, we altered the form of p_0_ by including a new constant repressive force that affects both normal and leukemic self-renewal resulting in 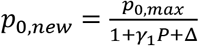 and 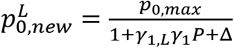 where *P* = *x*_*P*_*L*+ *x*_*P*_ and Δ is the differentiation therapy strength. We then performed combination therapy using our existing parameters, swept through differentiation therapy strengths and recorded which parameter sets achieved MR3 within 50 months. We then determined that a differentiation therapy strength of Δ=0.29 resulted in the highest proportion of parameter sets that achieved MR3 response. The full equations for combined TKI and differentiation therapy are shown in Supplemental Material **Eqns. (S40)-(S41)**.

### CML patient data

Data from newly diagnosed CML patients (n=21) treated with TKI therapy at UCI Health were obtained under an honest broker mechanism from the electronic health record, under Exemption 4 for human subjects research.

### Mice

C57BL/6J female mice (Jackson Laboratories), 6-12 weeks of age were used for irradiation and myeloid depletion experiments. Conditional *BCR-ABL1* double transgenic mice (*Koschmieder et al., 2005*) were obtained from Dr. Emmanuel Passegue (Columbia University). All protocols in mice were approved by Institutional Animal Use and Care Committee of University of California, Irvine.

### Irradiation of mice

To achieve selective depletion of hematopoietic stem cells (HSC), a 50cGy dose of irradiation from x-ray source (Precision X-rad 320) was applied. Control mice did not receive irradiation. The distribution of time points at which observations were made (days 1, 3, 7 post-irradiation), and the numbers of mouse replicates to use at each time point (between 2 and 7, totaling 13 mice), were informed by our Bayesian hierarchical framework for optimal experimental design (*Lomeli et al., 2021*).

### Myeloid cell depletion

RB6-8C5, an anti-Gr1 antibody (catalog # BE0075, BioXCell) or isotype control (catalog # BE0090, BioXCell) was injected intravenously, 50 μg per mouse, and mice sacrificed 24 hours later.

### BrdU injections

In irradiation experiments, mice were pulsed with BrdU by IP injection of 200 μl of 10 mg/ml BrdU in DPBS. BrdU flow kit (552598) from BD biosciences was used for detection of BrdU labeling in hematopoietic cells by flow cytometry.

### Flow cytometry analysis of cell populations

Bone marrow (BM) cells from femur and tibia of control and dosed mice were isolated by flushing bones. Following lysis of red blood cells (RBC lysis buffer, eBiosciences), leukocytes were stained with CD34 antibody for one hour and subsequently incubated with a cocktail of biotinylated antibodies directed against lineage markers (CD3, Gr-1, B220, Ter119) and stem/progenitor markers (c-Kit, Sca-1, CD48) for 30 minutes. Streptavidin (SA)-conjugated fluorchrome was utilized to detect biotynalated antibodies.

Following fixation, permeabilization, and DNase digestion, anti-BrdU antibody was used to assess BrdU incorporation. Events were acquired on FACS Arial II and analyzed with Flowjo v.10 software.

### Antibodies

Monoclonal antibodies for flow cytometry were biotinylated mouse lineage panel (559971, BD biosciences), PE-CF594 Streptavidin (562318, BD Biosciences), anti-mouse CD48 (561242, BD biosciences), anti-mouse CD34 eFluor450 (48-0341-82, eBiosciences), anti-mouse Sca-1-PE (108108, Biolegend), anti-mouse c-Kit-APC (17-1171-82, eBiosciences), FITC BrdU flow kit (559619, BD Biosciences).

### Generation and TKI treatment of chimeric *BCR-ABL1* mice

The full details of the CML mouse model will be published elsewhere (*Jena et al., 2022*). Briefly, bone marrow cells from conditional *BCR-ABL1* double transgenic mice (CD45.2^+^) (*Koschmieder et al., 2005*) (40 million cells) were transplanted intravenously into unirradiated C57BL/6J recipients (CD45.1^+^CD45.2^+^) maintained on doxycycline to suppress *BCR-ABL1* expression. Two months post-transplant, doxycycline was removed to allow induction of CML-like leukemia. Chimerism was assessed by percentage of CD45.1^−^ CD45.2^+^ granulocytes in peripheral blood. To generate chimeric mice with high (>90%) leukemic stem cell burden, the donor and recipient pair was reversed, with double transgenic mice transplanted with normal B6 BM. In mice with established CML-like leukemia (peripheral blood leukocytes > 20,000/μl and >40% circulating granulocytes), TKI treatment was initiated with dasatinib (25 mg/kg daily by oral gavage).

## Acknowledgements

The A.D.L., J.S.L. and R.A.V. acknowledge National Institutes of Health for partial support through grant nos. 1U54CA217378-01A1 for a National Center in Cancer Systems Biology at UC Irvine and P30CA062203 for the Chao Family Comprehensive Cancer Center at UC Irvine. In addition, A.D.L. and J.S.L. acknowledge support from DMS-1763272 and the Simons Foundation (594598QN) for an NSF-Simons Center for Multiscale Cell Fate Research, R.A.V. acknowledges support from R01 CA090576. J.S.L. also acknowledges NSF grants DMS-1936833 and DMS-1953410. J.R. and A.I. each acknowledge support from NSF graduate research fellowships.

## Author contributions

J.S.L., A.D.L., and R.A.V. designed the study. J.R., A.I., A.D.L. and J.S.L. developed the ordinary differential equation model. J.R. and A.I. developed the automated algorithm for the Design Space Analysis, performed the numerical simulations and analyzed the data. P.T., N.J., Z.-Y.L. and R.A.V. designed the mouse radiation and myeloid depletion experiments, which were performed by PT, and analyzed the experimental data. R.A.V. provided the anonymized CML patient data. J.R., A.I., P.T., A.D.L., J.S.L, and R.A.V. wrote the manuscript.

## Additional files

Supplemental Material

