## Supplemental Materials for "Predictive nonlinear modeling of malignant myelopoiesis and tyrosine kinase inhibitor therapy"

#### 1 Design Space Analysis

##### 1.1 Introduction

In this section, we provide details on the application of design space analysis (DSA), developed in [1, 2, 3] for chemical reaction networks, to a simplified version of the cell lineage model considered in the main text (see Fig. S2). This enables us to to analyze nonlinear dynamical systems near steady state to identify regions in parameter space where common qualitative behaviors occur. Applying this analysis allows us to (1) ignore specific parameter values, (2) obtain analytical steady states, (3) reduce the search area in parameter space by searching boundaries separating regions of common behaviors, and (4) easily automate this process. In this section, we review how one can construct a design space given an ordinary differential equation (ODE) model. We then apply this analysis to an ODE model of cell lineages and show how this model can be used to define regions of stability in parameter space.

##### 1.2 Boundaries of design space for general models

In order to apply DSA, the ODE must be a generalized mass action system, shown below in equations (1)- (2),

$$\frac{dX_i}{dt} = \sum_{k=1}^{r} \alpha_{ik} \prod_{j=1}^{n+m} X_j^{g_{ijk}} - \sum_{k=1}^{r} \beta_{ik} \prod_{j=1}^{n+m} X_j^{h_{ijk}}, \tag{1}$$

$$X_i(0) = X_{i0}, \tag{2}$$

for $i$=1,..,$n$. Here $n$ corresponds to the number of dependent variables and $m$ corresponds to the number of independent variables. The $\alpha_{ik}$ and $\beta_{ik}$ parameters correspond to rate constants of the differential equation and $r$ corresponds to the number of associated rate constants.

DSA takes advantage of the above form by creating a system of deconstructed ODEs where one source term and one sink term in the differential equation dominate, known as a sub-system (S-system). The following is the generalized form of a S-system:

$$\frac{dX_i}{dt} = \alpha_{ip} \prod_{j=1}^{n+m} X_j^{g_{ijp}} - \beta_{iq} \prod_{j=1}^{n+m} X_j^{h_{ijq}} \tag{3}$$

$$X_i(0) = X_{i0} \tag{4}$$

where $p$ and $q$ correspond to the number of positive and negative terms of the differential equation, respectively. We are interested in solving these solutions at steady state, and therefore solve the system by setting the time derivative to zero. We take advantage of the form shown in equation (3) and take the log of the system:

$$\log(\alpha_{ip}) + \sum_{j=1}^{n+m} g_{ijp} \log(X_j) = \log(\beta_{iq}) + \sum_{j=1}^{n+m} h_{ijq} \log(X_j) \tag{5}$$

thus, making this a linear solve in log space. In defining the S-systems, we must make assumptions about the model and its parameters. To satisfy the S-systems, we impose inequality constraints to satisfy the dominating source and sink terms of the S-systems by the following,

$$\alpha_{ip} \prod_{j=1}^{n+m} X_j^{g_{ijp}} > \alpha_{i\bar{p}} \prod_{j=1}^{n+m} X_j^{g_{ij\bar{p}}} \text{for i = 1,..,n; } \bar{p}\text{=1,...,p-1,p+1,...,r} \tag{6}$$

$$\beta_{iq} \prod_{j=1}^{n+m} X_j^{h_{ijq}} > \beta_{i\bar{q}} \prod_{j=1}^{n+m} X_j^{h_{ij\bar{q}}} \text{for i = 1,..,n; } \bar{q}\text{=1,...,q-1,q+1,...,r} \tag{7}$$

We can then log transform these inequalities to obtain

$$\log(\alpha_{ip}) + \sum_{j=1}^{n+m} g_{ijp} \log(X_j) > \log(\alpha_{i\bar{p}}) + \sum_{j=1}^{n+m} g_{ij\bar{p}} \log(X_j) \tag{8}$$

$$\log(\beta_{iq}) + \sum_{j=1}^{n+m} h_{ijq} \log(X_j) > \log(\beta_{i\bar{q}}) + \sum_{j=1}^{n+m} h_{ij\bar{q}} \log(X_j) \tag{9}$$

These inequalities become our boundaries in parameter space in which qualitative behavior is shared once the $X_j$'s are evaluated at steady state, where the steady states are obtained from solving the log-linear S-system. Not all S-systems will have a unique solution or satisfy the inequality constraints. The S-systems that do not satisfy these constraints will be discarded. The analysis is summarized in figure S1.

#### 1.3 Analysis of a four cell lineage model

We next apply this analysis to a lineage model with four cell types (see Fig. S2). The model consists of two dividing cell types consisting of $S$ (HSC) and $P$ (MPP) cells with a division rate associated to the cells ($\eta_1$ and $\eta_2$ respectively). The $S$ cells have the ability to self renew with fraction ($p_0$) or differentiate ($1 - p_0$). The $P$ cells have the ability to self renew with fraction ($p_1$) or differentiate into either $TD_l$ (lymphoid) or $TD_m$ (myeloid) cells ($q_1$ or $1-p_1-q_1$ respectively). $TD_m$ and $TD_l$ cells only have the ability to die at rates $d_m$ and $d_l$, respectively. We add negative feedback on the self renewal fraction of the stem cells from the differentiated cells. Figure S2 shows a schematic of the lineage with the parameters. The corresponding differential equations are

$$x'_S = (2p_0 - 1)\eta_1 x_S \tag{10}$$
$$x'_P = 2(1 - p_0)\eta_1 x_S + (2p_1 - 1)\eta_2 x_P \tag{11}$$
$$x'_{TD_L} = 2q_1\eta_2 x_P - d_l x_{TD_l} \tag{12}$$
$$x'_{TD_M} = 2(1 - p_1 - q_1)\eta_2 x_P - d_m x_{TD_m} \tag{13}$$

where $p_0 = \bar{p}_0/(1 + \gamma x_P)$ and $\bar{p_0}$ is defined as the maximum stem cell self renewal fraction and $\gamma$ is the feedback strength.

We begin by rewriting the equations in the form shown in equation (1).

$$x'_S = (2\bar{p}_0 x_{new}^{-1} - 1)\eta_1 x_S \tag{14}$$
$$x'_P = 2(1 - \bar{p}_0 x_{new}^{-1})\eta_1 x_S + (2p_1 - 1)\eta_2 x_P \tag{15}$$
$$x'_{TD_L} = 2q_1\eta_2 x_P - d_l x_{TD_l} \tag{16}$$
$$x'_{TD_M} = 2(1 - p_1 - q_1)\eta_2 x_P - d_m x_{TD_m} \tag{17}$$
$$0 = 1 + \gamma x_P - x_{new}. \tag{18}$$

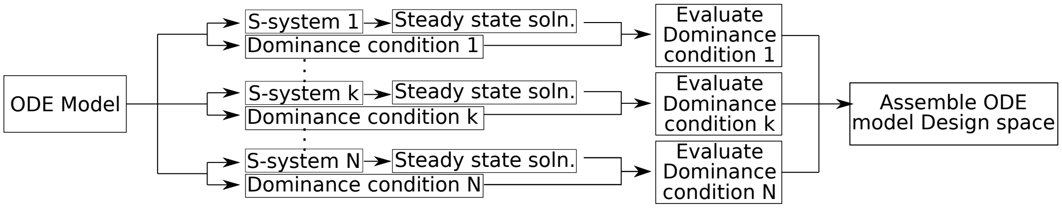


Figure S1: Flow chart for DSA. Given an ODE, we can obtain the design space by obtaining all S-systems, steady states, and evaluated dominance conditions. S-systems that do not satisfy the dominance condition are not included in the design space.

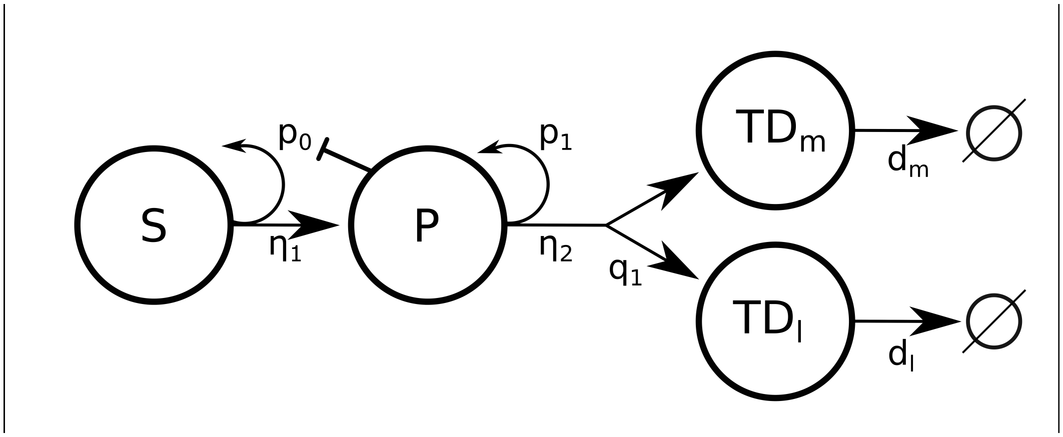


Figure S2: Lineage schematic depicting a stem cell and terminally differentiated cell with negative feedback onto stem cell self-renewal probability $p_0$.

Note, we introduce a new variable $x_{new} = 1 + \gamma x_P$, which is necessary to achieve the form of equation (1). We next find all combinations in which one source term and one sink term dominates the system of differential equations.

From equations (14)-(18), we obtain 24 S-System combinations, a subset of which is shown in table S1. We continue the analysis with S-system 2 from table S1. Using S-system 2, we set the time derivatives to zero and rearrange the equations such that we obtain $A\bar{x} = \bar{b}$:

| | S-system 1 | S-system 2 | S-system 3 | S-system 4 |
|---|---|---|---|---|
| $x'_S$ | $2\bar{p_0}x_{new}^{-1}\eta_1 x_S - \eta_1 x_S$ | $2\bar{p_0}x_{new}^{-1}\eta_1 x_S - \eta_1 x_S$ | $2\bar{p_0}x_{new}^{-1}\eta_1 x_S - \eta_1 x_S$ | $2\bar{p_0}x_{new}^{-1}\eta_1 x_S - \eta_1 x_S$ |
| $x'_P$ | $2\eta_1 x_S - 2\bar{p_0}x_{new}^{-1}\eta_1 x_S$ | $2\eta_1 x_S - \eta_2 x_S$ | $2p_1\eta_2 x_S - 2\bar{p_0}x_{new}^{-1}\eta_1 x_S$ | $2p_1\eta_2 x_S - \eta_2 x_S$ |
| $x'_{TD_l}$ | $2q_1\eta_2 x_P - d_l x_{TD_l}$ | $2q_1\eta_2 x_P - d_l x_{TD_l}$ | $2q_1\eta_2 x_P - d_l x_{TD_l}$ | $2q_1\eta_2 x_P - d_l x_{TD_l}$ |
| $x'_{TD_m}$ | $2\eta_2 x_P - d_m x_{TD_m}$ | $2\eta_2 x_P - d_m x_{TD_m}$ | $2\eta_2 x_P - d_m x_{TD_m}$ | $2\eta_2 x_P - d_m x_{TD_m}$ |
| 0 | $\gamma x_P - x_{new}$ | $\gamma x_P - x_{new}$ | $\gamma x_P - x_{new}$ | $\gamma x_P - x_{new}$ |
| Boundary 1 | $2\eta_1 x_S > 2p_1\eta_2 x_S$ | $2\eta_1 x_S > 2p_1\eta_2 x_S$ | $2\eta_1 x_S < 2p_1\eta_2 x_S$ | $2\eta_1 x_S < 2p_1\eta_2 x_S$ |
| Boundary 2 | $2\bar{p_0}x_{new}^{-1}\eta_1 x_S > \eta_2 x_S$ | $2\bar{p_0}x_{new}^{-1}\eta_1 x_S < \eta_2 x_S$ | $2\bar{p_0}x_{new}^{-1}\eta_1 x_S > \eta_2 x_S$ | $2\bar{p_0}x_{new}^{-1}\eta_1 x_S < \eta_2 x_S$ |

Table S1: A sample of S-systems from equations (14)-(18).

$$\begin{bmatrix} 0 & 0 & 0 & 0 & 1 \\ 1 & -1 & 0 & 0 & 0 \\ 0 & 1 & -1 & 0 & 0 \\ 0 & 1 & 0 & -1 & 0 \\ 0 & -1 & 0 & 0 & 1 \end{bmatrix} \begin{bmatrix} \log(\bar{x_S}) \\ \log(\bar{x_P}) \\ \log(\bar{x}_{TD_l}) \\ \log(\bar{x}_{TD_m}) \\ \log(\bar{x}_{new}) \end{bmatrix} = \begin{bmatrix} \log(2\bar{p_0}) \\ \log(\frac{\eta_2}{2\eta_1}) \\ \log(\frac{d_l}{2q_1\eta_2}) \\ \log(\frac{d_m}{2\eta_2}) \\ \log(\gamma) \end{bmatrix}$$

such that $\bar{x_S}$, $\bar{x_P}$, $\bar{x}_{TD_l}$, and $\bar{x}_{TD_m}$ are the steady state solutions for the S-system and $\bar{x}_{new}$ is the solution of the newly defined variable at steady state. Solving the linear equation gives us:

$$\begin{bmatrix} \log(\bar{x_S}) \\ \log(\bar{x_P}) \\ \log(\bar{x}_{TD_l}) \\ \log(\bar{x}_{TD_m}) \\ \log(\bar{x}_{new}) \end{bmatrix} = \begin{bmatrix} \log(\frac{\bar{p_0}\eta_2}{\gamma}) \\ \log(\frac{2\bar{p_0}}{\gamma}) \\ \log(\frac{4\bar{p_0}q_1\eta_2}{d_l}) \\ \log(\frac{4\bar{p_0}\eta_2}{d_m}) \\ \log(2\bar{p_0}) \end{bmatrix}.$$

We can now construct the boundaries for this S-System. We substitute our steady state solutions obtained above into the logged inequality constraints from table S1. Thus, the inequalities in log space become

$$0 < \log(2\bar{p_0}) \quad (19)$$
$$0 > \log(2\bar{p_1}) \quad (20)$$
$$0 > \log(\bar{q_1}) \quad (21)$$

Out of the 24 possible S-systems, only S-system 2 has a unique steady state that satisfies the constraints. We can plot the design space by varying $\bar{p_0}$ and $\gamma$ (see Fig. S3). The design space shows one region where $\bar{p_0} > 0.5$, a requirement for a positive steady state in the full system. The domain in parameter space corresponds to S-system 2 in table S1.

It is possible to relate the S-system back to the true ODE system with classical techniques. For example, it was shown in [1, 2, 3] that the S-system and full ODE system have the same linear stability behavior in the parameter regime appropriate for the S-system. Thus, parameter sensitivity analyses of the S-system provide insight on the behavior of the full system.

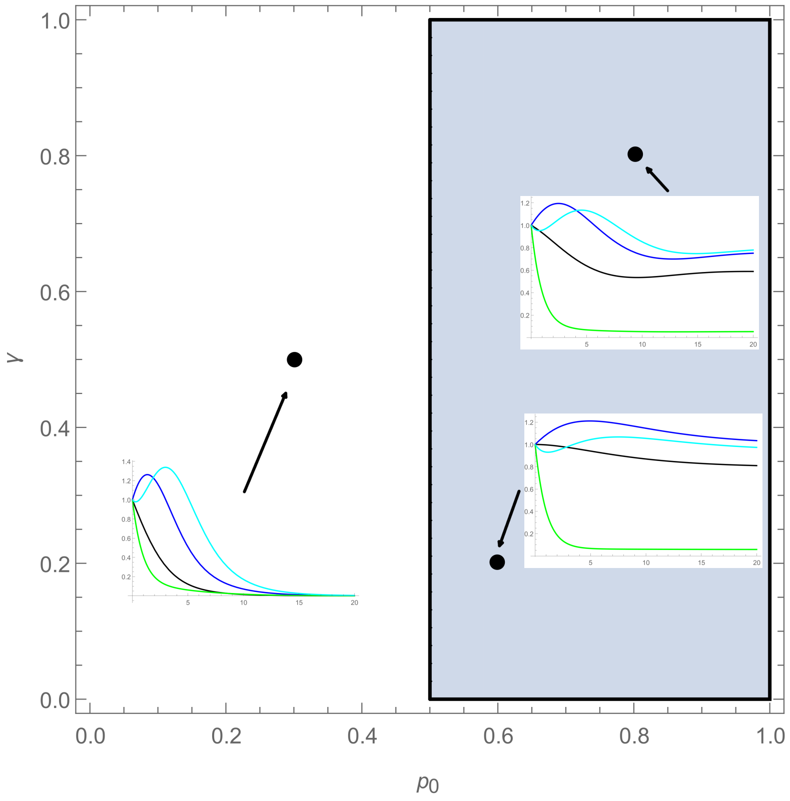


Figure S3: Design space for four cell lineage model. The design space is showing a slice of parameter space varying the max self renewal probability ($\bar{p_0}$) and feedback gain for stem cells ($\gamma$). If we are to sample parameters sets (shown by points), we observe oscillatory behavior in the full system, as shown by the time evolution plots.

Performing a linear stability analysis on the S-system that corresponds to the region in parameter space shown in figure S3, we obtain the following eigenvalues:

$$\lambda = [-0.5d - \frac{0.125\gamma\sqrt{(d(-64+16d)\bar{p_0}^4)/\gamma^2}}{\bar{p_0}^2}, \tag{22}$$

$$-0.5d + \frac{0.125\gamma\sqrt{(d(-64+16d)\bar{p_0}^4)/\gamma^2}}{\bar{p_0}^2}] \tag{23}$$

which suggests a stable spiral or stable node depending on the value of d. We also perform the linear stability analysis on the full system and obtain the following eigenvalues:

$$\lambda = [\frac{-0.25d\bar{p_0} - 0.125\sqrt{d\bar{p_0}^2(4d+(32-64p_0)\bar{p_0})}}{\bar{p_0}^2}, \tag{24}$$

$$\frac{-0.25dp_0 + 0.125\sqrt{d\bar{p_0}^2(4d+(32-64\bar{p_0})\bar{p_0})}}{\bar{p_0}^2}] \tag{25}$$

and in the parameter space for the S-system, we obtain the same behavior. Using the full system, we can also plot the dynamics in the domain, which are shown as insets in figure S3. When plotting the full system in figure S3, we observe that the stem and progenitor populations oscillate before reaching steady state, characteristics of a stable spiral. The S-system and the true system eigenvalues are not identical, but using the same parameters in the appropriate S-system parameter space, the systems will have the same behavior. However, outside of the domain of validity of the S-system, we cannot conclude anything about the full system's dynamics from the S-system. In fact, from figure S3, we observe being outside of the domain of validity of S-system 2 (e.g.,$\bar{p}_0 < 0.5$) yields to solutions that tend to the zero steady state.

We use DSA to select among all possible ODE models for normal hematopoiesis consistent with the lineage diagram shown in Fig. 1A in the main text in which each parameter $p_0$, $p_1$, $q_1$, $\eta_1$, and $\eta_2$ is either unregulated (constant) or is subject to positive or negative regulation from most one cell type in the lineage. As described in the main text, there are 59,049 possible models, counting each combination as a model. We implement an automated implementation of DSA that enables an efficient exploration of this large space of models. Eliminating models with no valid S-systems and those with unstable equilibria, we eliminate all but the four model classes shown in Fig. 1B in the main text. All the models have negative regulation of the HSCs but differ in the cell type that provides the regulation. The models within the classes share at least one S-system; the differences between models in a class are the types of regulation (positive, negative, none) on the other parameters and the cells that provide the regulation.

#### 2 Mathematical Model

The complete ODE model of normal hematopoiesis is composed of the following equations

$$x'_S = \left(2\frac{p_{0,max}}{1+\gamma_1 x_P} - 1\right)\frac{\eta_{1,max}}{1+\gamma_2 x_S} x_S \tag{26}$$

$$x'_P = 2\left(1 - \frac{p_{0,max}}{1+\gamma_1 x_P}\right)\frac{\eta_{1,max}}{1+\gamma_2 x_S} x_S + \left(2\frac{p_{1,max}}{1+\gamma_3 x_{TD_m}} - 1\right)\frac{\eta_{2,max}}{1+\gamma_5 x_S} x_P \tag{27}$$

$$x'_{TD_L} = 2\frac{q_{1,max}}{1+\gamma_4 x_{TD_m}}\frac{\eta_{2,max}}{1+\gamma_5 x_S} x_P - d_L x_{TD_L} \tag{28}$$

$$x'_{TD_m} = 2\left(1 - \frac{p_{1,max}}{1+\gamma_3 x_{TD_m}} - \frac{q_{1,max}}{1+\gamma_4 x_{TD_m}}\right)\frac{\eta_{2,max}}{1+\gamma_5 x_S} x_P - d_m x_{TD_m}. \tag{29}$$

The ODE model for CML hematopoiesis tracks the dynamics of both the normal and CML cells (superscript $L$), and assumes that both cell types provide and respond to feedback signaling, although the CML stem cells are slightly less responsive to negative feedback regulation, which gives them a fitness advantage. The complete system is given by

$$x'_S = \left(2\frac{p_{0,max}}{1+\gamma_1(x_P + x_{P^L})} - 1\right)\frac{\eta_{1,max}}{1+\gamma_2(x_S + x_{S^L})}x_S \tag{30}$$

$$x'_P = 2\left(1 - \frac{p_{0,max}}{1+\gamma_1(x_P + x_{P^L})}\right)\frac{\eta_{1,max}}{1+\gamma_2(x_S + x_{S^L})}x_S \tag{31}$$
$$+ \left(2\frac{p_{1,max}}{1+\gamma_3(x_{TD_m} + x_{TD_m^L})} - 1\right)\frac{\eta_{2,max}}{1+\gamma_5(x_S + x_{S^L})}x_P$$

$$x'_{TD_L} = 2\frac{q_{1,max}}{1+\gamma_4(x_{TD_m} + x_{TD_m^L})}\frac{\eta_{2,max}}{1+\gamma_5(x_S + x_{S^L})}x_P - d_L x_{TD_L} \tag{32}$$

$$x'_{TD_m} = 2\left(1 - \frac{p_{1,max}}{1+\gamma_3(x_{TD_m} + x_{TD_m^L})} - \frac{q_{1,max}}{1+\gamma_4(x_{TD_m} + x_{TD_m^L})}\right)\frac{\eta_{2,max}}{1+\gamma_5(x_S + x_{S^L})}x_P$$
$$- d_m x_{TD_m} \tag{33}$$

$$x'_{S^L} = \left(2\frac{p_{0,max}}{1+\gamma_{1,L}\gamma_1(x_P + x_{P^L})} - 1\right)\frac{\eta_{1,max}}{1+\gamma_{2,L}\gamma_2(x_S + x_{S^L})}x_{S^L} \tag{34}$$

$$x'_{P^L} = 2\left(1 - \frac{p_{0,max}}{1+\gamma_{1,L}\gamma_1(x_P + x_{P^L})}\right)\frac{\eta_{1,max}}{1+\gamma_{2,L}\gamma_2(x_S + x_{S^L})}x_{S^L}$$
$$+ \left(2\frac{p_{1,max}}{1+\gamma_{3,L}\gamma_3(x_{TD_m} + x_{TD_m^L})} - 1\right)\frac{\eta_{2,max}}{1+\gamma_{5,L}\gamma_5(x_S + x_{S^L})}x_{P^L} \tag{35}$$

$$x'_{TD_L^L} = 2\frac{q_{1,max}}{1+\gamma_{4,L}\gamma_4(x_{TD_m} + x_{TD_m^L})}\frac{\eta_{2,max}}{1+\gamma_{5,L}\gamma_5(x_S + x_{S^L})}x_{P^L} - d_L x_{TD_L^L} \tag{36}$$

$$x'_{TD_m^L} = 2\left(1 - \frac{p_{1,max}}{1+\gamma_{3,L}\gamma_3(x_{TD_m} + x_{TD_m^L})} - \frac{q_{1,max}}{1+\gamma_{4,L}\gamma_4(x_{TD_m} + x_{TD_m^L})}\right)\frac{\eta_{2,max}}{1+\gamma_{5,L}\gamma_5(x_S + x_{S^L})}x_{P^L}$$
$$- d_m x_{TD_m^L}. \tag{37}$$

When TKI therapy is applied, equations (34) and (35) are replaced with

$$x'_{S^L} = \left(2\frac{p_{0,max}}{1+\gamma_{1,L}\gamma_1(x_P + x_{P^L})} - 1 - TKI_{HSC}\right)\frac{\eta_{1,max}}{1+\gamma_{2,L}\gamma_2(x_S + x_{S^L})}x_{S^L} \tag{38}$$

$$x'_{P^L} = 2\left(1 - \frac{p_{0,max}}{1+\gamma_{1,L}\gamma_1(x_P + x_{P^L})}\right)\frac{\eta_{1,max}}{1+\gamma_{2,L}\gamma_2(x_S + x_{S^L})}x_{S^L}$$
$$+ \left(2\frac{p_{1,max}}{1+\gamma_{3,L}\gamma_3(x_{TD_m} + x_{TD_m^L})} - 1 - TKI_{MPP}\right)\frac{\eta_{2,max}}{1+\gamma_{5,L}\gamma_5(x_S + x_{S^L})}x_{P^L}, \tag{39}$$

where $TKI_{HSC}$ and $TKI_{MPP}$ denote TKI-induced death rates of the CML stem and MPP cells. With the introduction of differentiation therapy, eqns. (30) and (38) instead become

$$x'_S = \left(2\frac{p_{0,max}}{1+\gamma_1(x_P + x_{P^L}) + \Delta} - 1\right)\frac{\eta_{1,max}}{1+\gamma_2(x_S + x_{S^L})}x_S \tag{40}$$

$$x'_{S^L} = \left(2\frac{p_{0,max}}{1+\gamma_{1,L}\gamma_1(x_P + x_{P^L}) + \Delta} - 1 - TKI_{HSC}\right)\frac{\eta_{1,max}}{1+\gamma_{2,L}\gamma_2(x_S + x_{S^L})}x_{S^L}. \tag{41}$$

### 3 Parameter estimation

The parameters for the model of normal hematopoesis, and their descriptions, are listed below in Table S2. The additional parameters needed to model the CML cell population, and the application of TKI therapy are given in Table S3.

| Parameter | Description |
|---|---|
| $p_{0,max}$ | Maximal self-renewal fraction of HSC |
| $p_{1,max}$ | Maximal self-renewal fraction of MPP |
| $q_{1,max}$ | Maximum branching fraction from MPP to $TD_L$ |
| $\eta_{1,max}$ | maximum HSC proliferation rate |
| $\eta_{2,max}$ | maximum MPP proliferation rate |
| $\gamma_1$ | Feedback gain on HSC self-renewal fraction |
| $\gamma_2$ | Feedback gain on HSC proliferation rate (from HSC) |
| $\gamma_3$ | Feedback gain on MPP self-renewal fraction |
| $\gamma_4$ | Feedback gain on MPP branching fraction |
| $\gamma_5$ | Feedforward gain on MPP proliferation rate |
| $d_L$ | Death rate of $TD_L$ |
| $d_m$ | Death rate of $TD_M$ |

Table S2: Parameter values used to model the normal hematopoietic system.

| Parameter | Description |
|---|---|
| $\gamma_{1,L}$ | Modified feedback gain on HSC self-renewal fraction |
| $\gamma_{2,L}$ | Modified feedback gain on HSC proliferation |
| $\gamma_{3,L}$ | Modified feedback gain on MPP self-renewal fraction |
| $\gamma_{4,L}$ | Modified feedback gain on MPP branching fraction |
| $\gamma_{5,L}$ | Modified feedforward gain on MPP proliferation |
| $TKI_{HSC}$ | Death rate of HSC due to TKI therapy |
| $TKI_{MPP}$ | Death rate of MPP due to TKI therapy |

Table S3: Additional parameter values used to model the CML cell population dynamics and effect of TKI therapy.

#### 3.1 Parameter distributions

A grid-search algorithm was to find parameter values that demonstrate steady state cell counts consistent with [4] and time of recovery to steady state from cellular perturbations consistent with [5]. The resulting distributions of the 1493 parameters are displayed along the diagonal of figure S4. In the distributions the feedback gain of the feed-forward regulation is skewed towards smaller values across all parameter sets. To investigate the observed feed-forward loop, we selected only the parameter sets with sufficiently large feedforward gain $\gamma_5 > 0.01$. This resulted in a reduced distribution of 563 parameter sets and is shown in S5. The parameter ranges for the gridsearch were found using the method shown in the following pseudo-Python.

```
for i in range(1e6):
```

```
param_gridsearch_dist = dict(p0max=np.random.uniform(0.5, 1.0),
p1max=np.random.uniform(0.0,0.5), q1max=np.abs(np.random.uniform(0.0,0.49)),
eta1max=np.random.uniform(0,0.5), eta2max=10**np.random.uniform(-2,1.5),
gam2=10**np.random.uniform(-6,0), gam3=10**np.random.uniform(-6,0),
gam4=10**np.random.uniform(-6,0), gam5=10**np.random.uniform(-6,0),
dL=10**np.random.uniform(-4,1), dM=10**np.random.uniform(-4,1))

y = hematopoiesis(param_gridsearch_dist)

if y[0,-1]>.01 and y[0,-1]<y[1,-1]<y[2,-1]<y[3,-1]:
    master_params.append(param_gridsearch_dist)
```

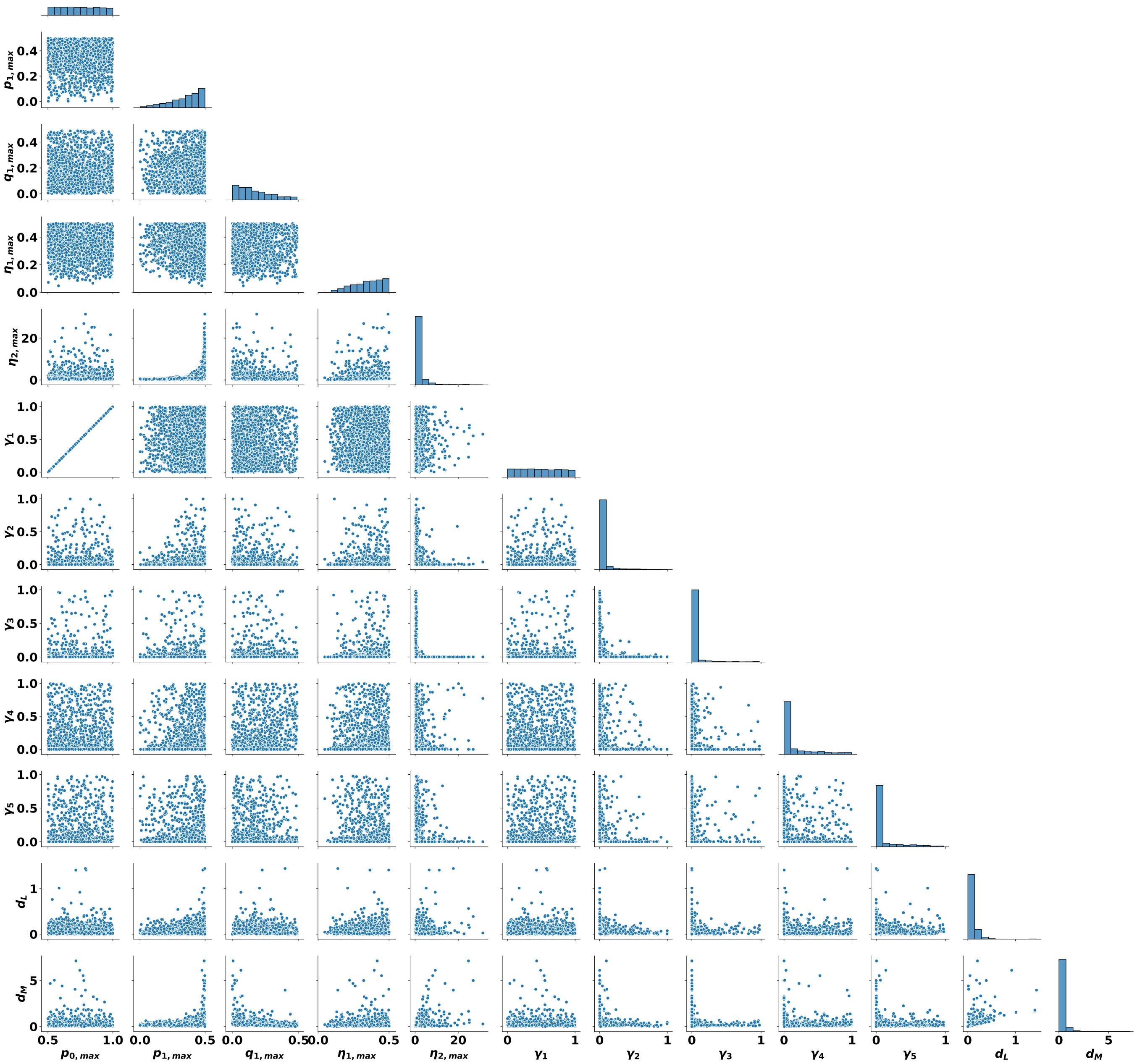


Figure S4: Pairwise parameter distributions for all 1493 parameter sets found from the grid-search. The overall distributions for each parameter are shown along the diagonal.

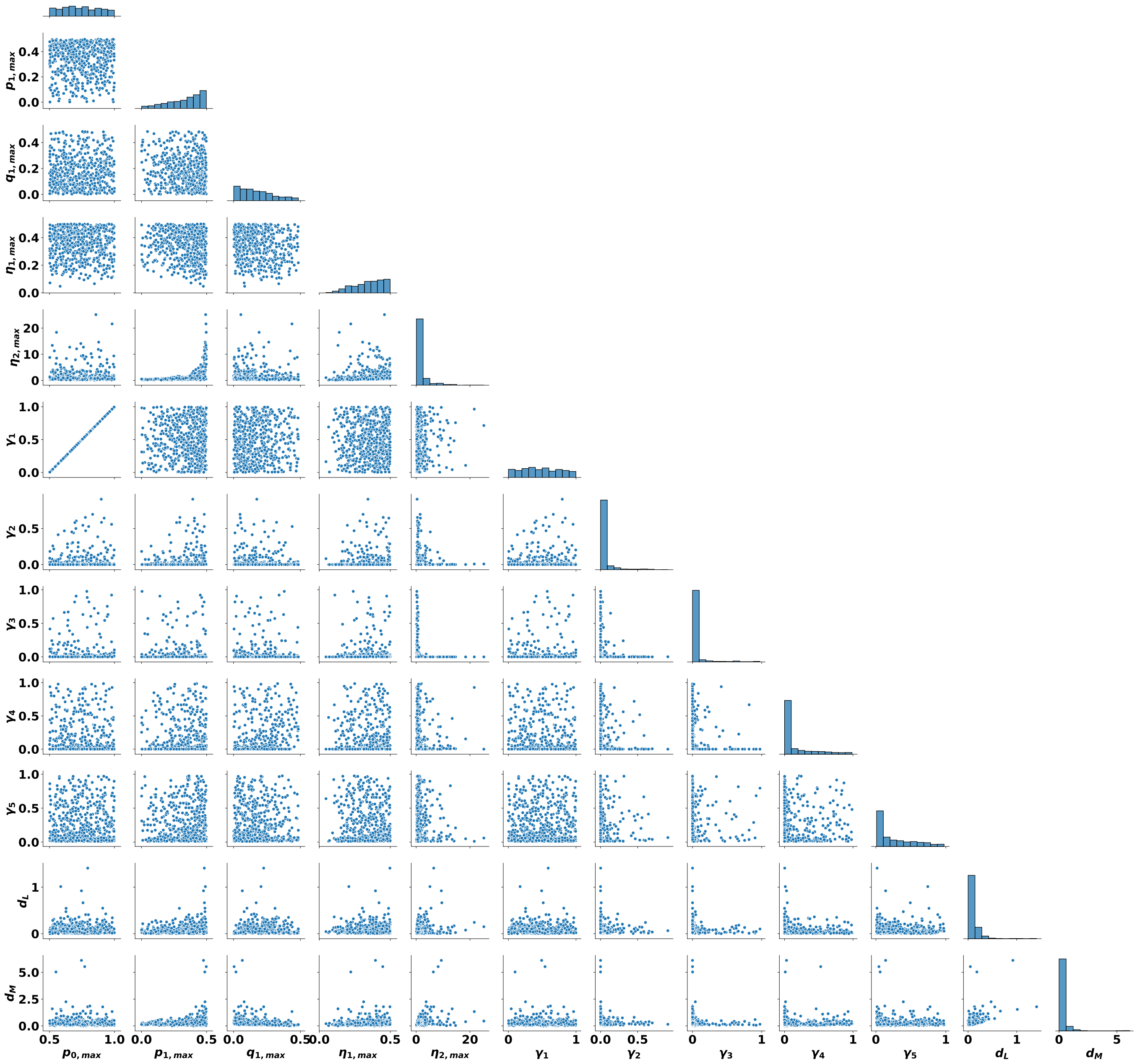


Figure S5: Pairwise parameter distributions for the 563 parameter sets from Fig. S4 that have feedforward gain $\gamma_5 > 0.01$. The overall distributions for each parameter are shown along the diagonal.

The values of the parameters used to model the normal hematopoietic system in Figs. 3-6 in the main text are given in Table S4. When CML cells are introduced, the values of the additional parameters associated with the leukemic cell ODEs are given in Table S5.

| Parameter | Value |
|---|---|
| $p_{0,max}$ | 0.756641 |
| $p_{1,max}$ | 0.357913 |
| $q_{1,max}$ | 0.032241 |
| $\eta_{1,max}$ | 0.197639 |
| $\eta_{2,max}$ | 0.470880 |
| $\gamma_1$ | 0.513281 |
| $\gamma_2$ | 0.543987 |
| $\gamma_3$ | 0.000165 |
| $\gamma_4$ | 0.000428 |
| $\gamma_5$ | 0.201311 |
| $d_L$ | 0.024757 |
| $d_m$ | 0.374514 |

Table S4: Parameter values used to model the normal hematopoietic system in Figs. 3C-6 in the main text. In Figs. 3A and 3B, in region *i*: $\gamma_1 = 0.095$, $\gamma_3 = 0.3$; in region *ii*: $\gamma_1 = 2.0$, $\gamma_3 = 1.0$, in region *iii* the parameters listed in the table are used. All other parameters are the same as listed in the table.

| Parameter | Value |
|---|---|
| $\gamma_{1,L}$ | 0.5 |
| $\gamma_{2,L}$ | 1 |
| $\gamma_{3,L}$ | 1 |
| $\gamma_{4,L}$ | 1 |
| $\gamma_{5,L}$ | 1 |
| $TKI_{HSC}$ | 0.25 |
| $TKI_{MPP}$ | 30 |

Table S5: Additional parameter values used to model the CML cell population dynamics.

### 4 Sensitivity analysis

Here we present the results of sensitivity analyses for the normal and CML hematopoiesis models. See captions of the figures for details.

#### 4.1 Results of individual perturbations for normal hematopoiesis model

| Parameter | Value |
|---|---|
| $p_{0,max}$ | 0.57566 |
| $p_{1,max}$ | 0.025144 |
| $q_{1,max}$ | 0.341495 |
| $\eta_{1,max}$ | 0.338547 |
| $\eta_{2,max}$ | 0.346647 |
| $\gamma_1$ | 0.15132 |
| $\gamma_2$ | 0.000251023 |
| $\gamma_3$ | 0.345963 |
| $\gamma_4$ | 2.87419e-06 |
| $\gamma_5$ | 0.964583 |
| $d_L$ | 0.0585988 |
| $d_m$ | 0.0857891 |

Table S6: Parameter values used to model a representative non-responsive individual in Figs. 6-7 in the main text with leukemic parameters from Table S5.

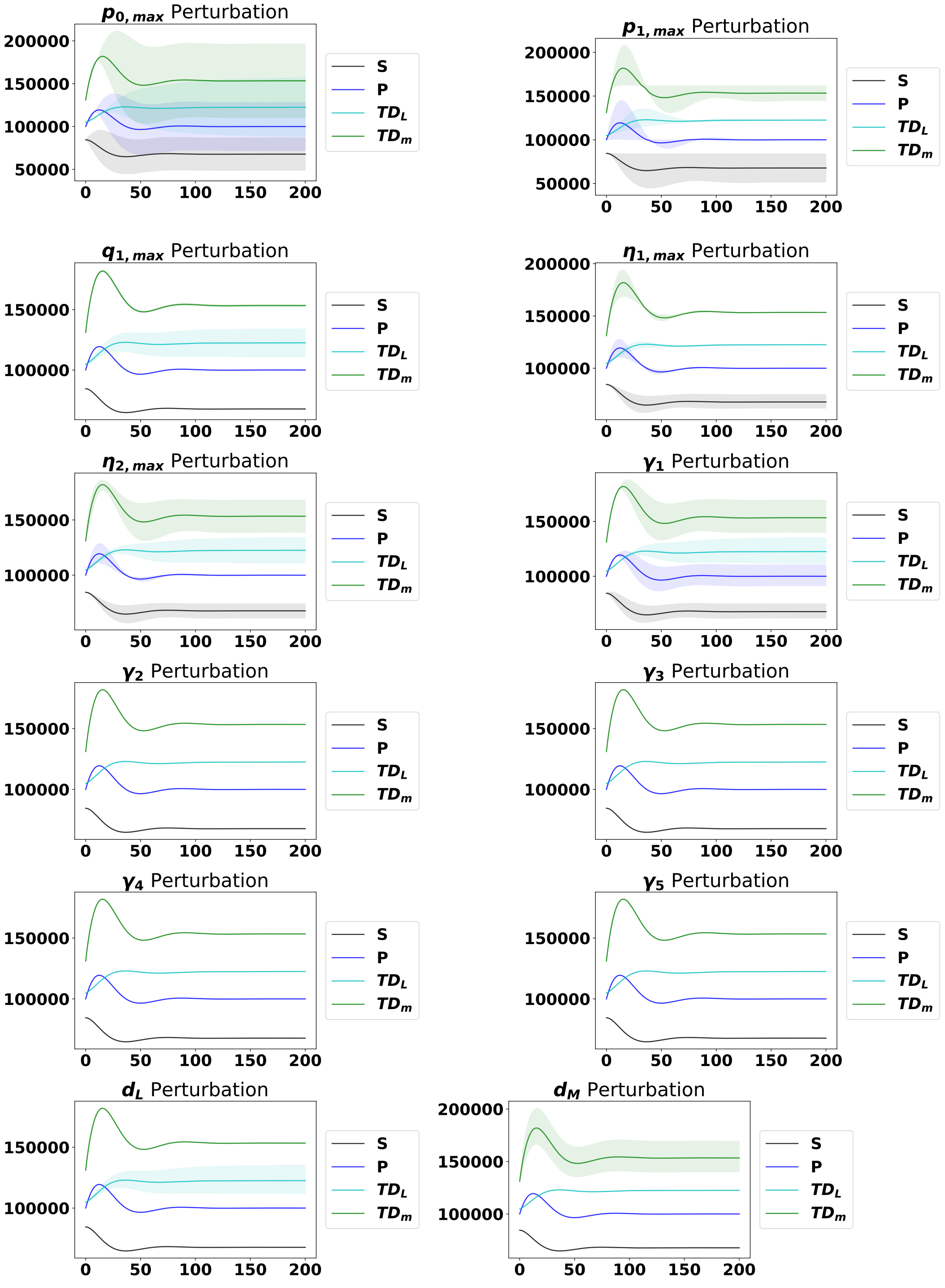


Figure S6: The dynamics from the steady state with perturbations of each parameter, as labeled, ranging from 90% to 110% of the original parameter value. The solid line represents the median of the perturbations, while the shaded region represents 95% of the perturbation dynamics.

#### 4.2 Results of individual perturbations for CML hematopoiesis model

**A**

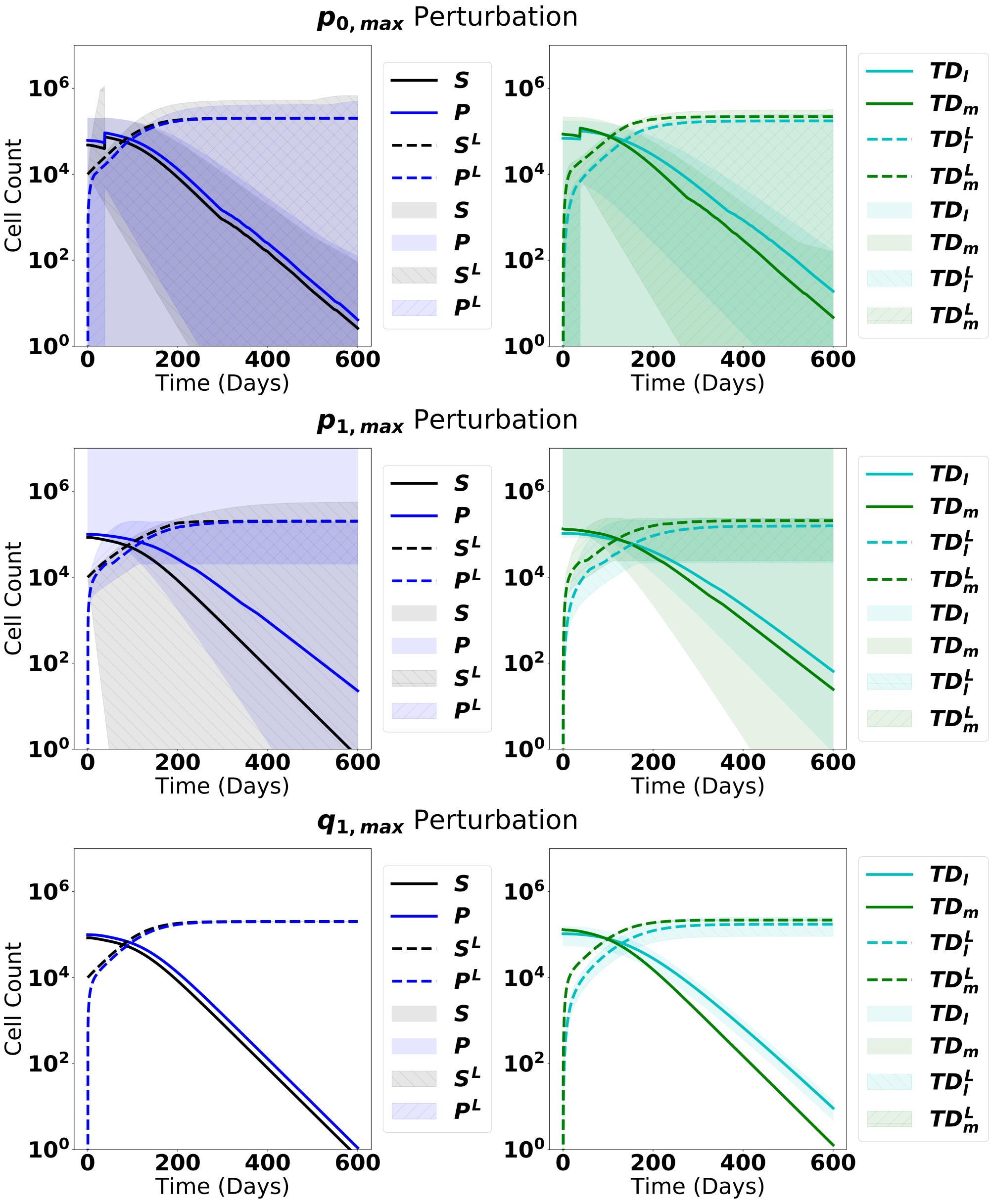

**B**

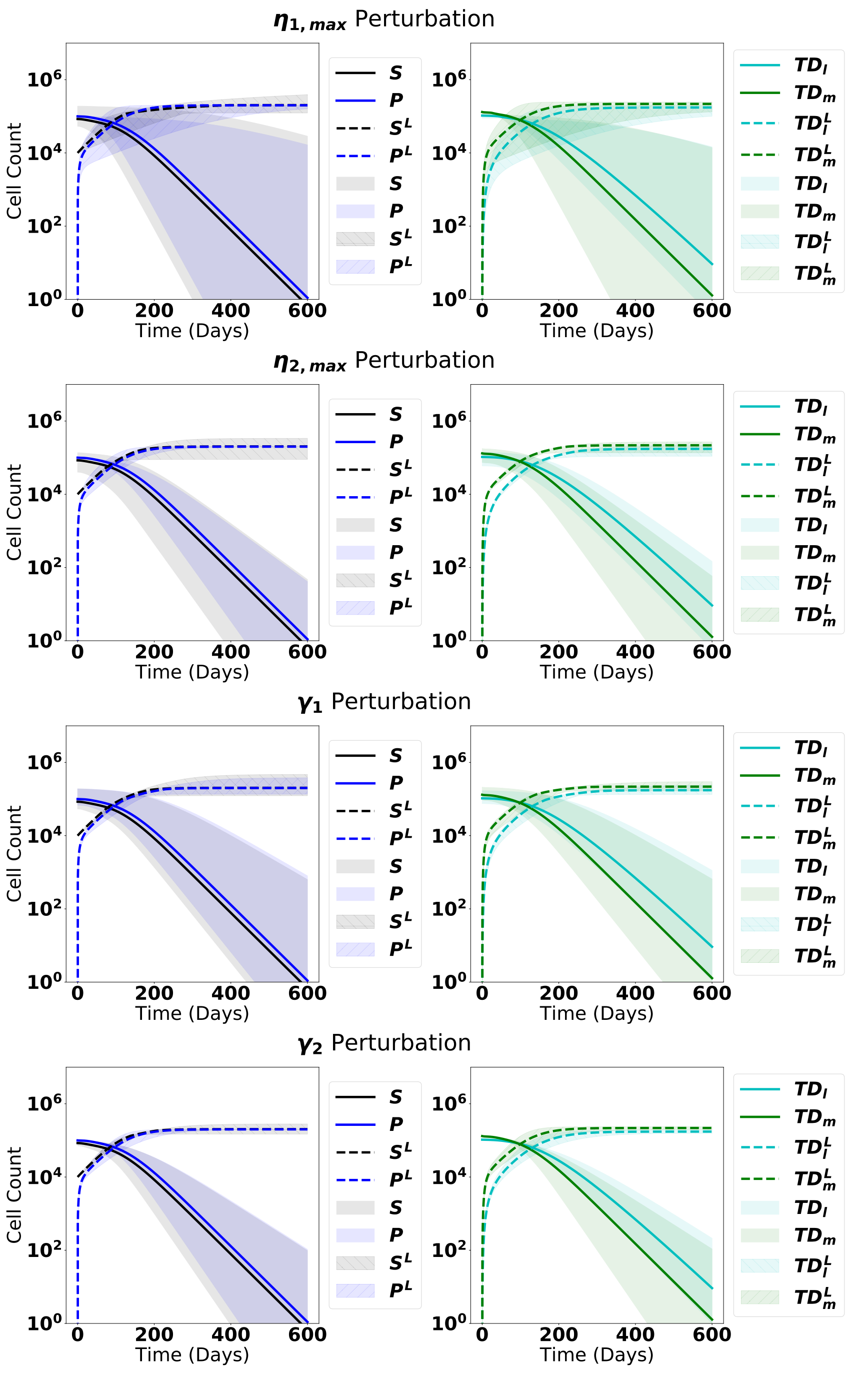
$\eta_{1,max}$ Perturbation
Cell Count
$10^0$ $10^2$ $10^4$ $10^6$
0 200 400 600
Time (Days)
$S$ $P$ $S^L$ $P^L$ $S$ $P$ $S^L$ $P^L$
$TD_l$ $TD_m$ $TD_l^L$ $TD_m^L$ $TD_l$ $TD_m$ $TD_l^L$ $TD_m^L$
$\eta_{2,max}$ Perturbation
Cell Count
Time (Days)
$\gamma_1$ Perturbation
Cell Count
Time (Days)
$\gamma_2$ Perturbation
Cell Count
Time (Days)

**C**

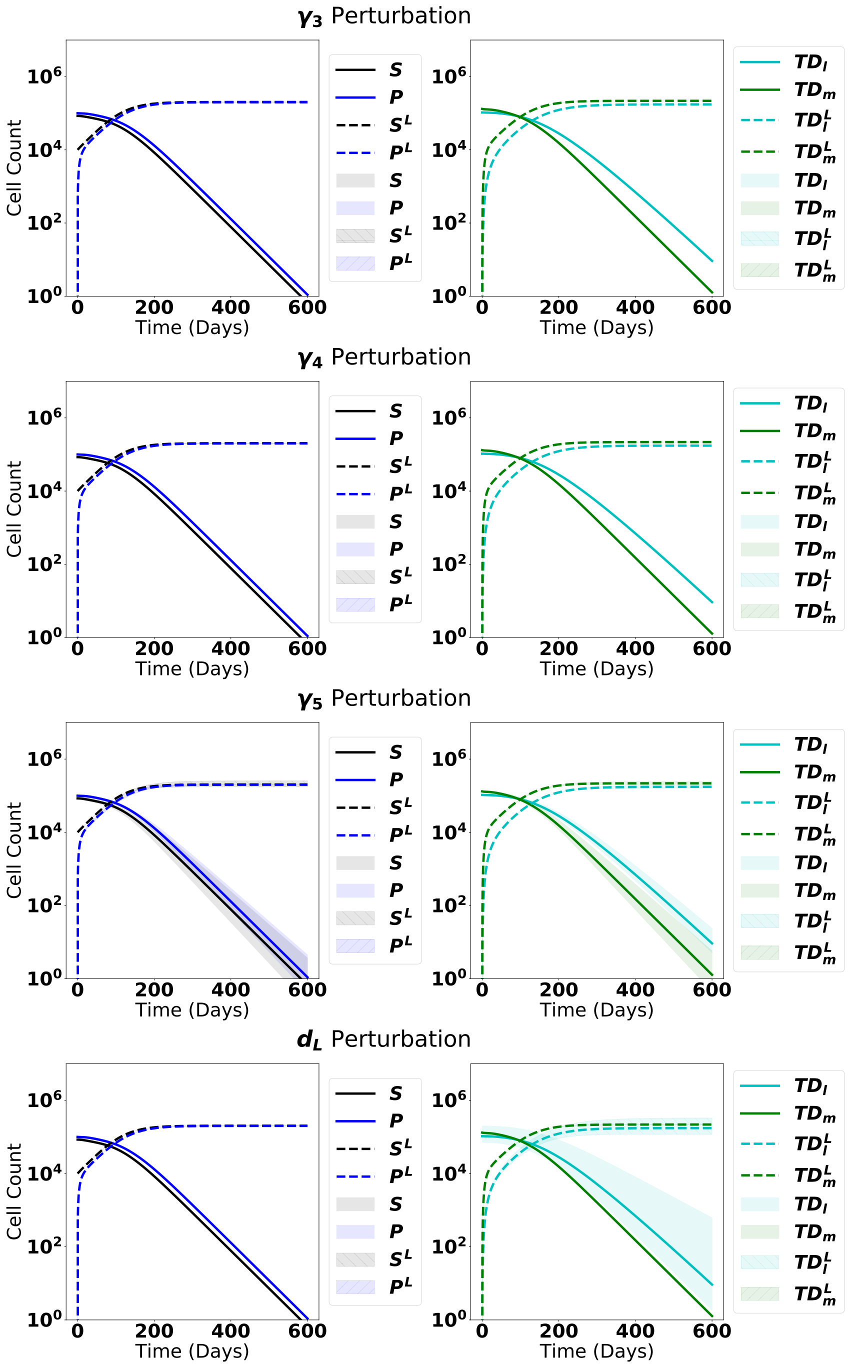

γ3 Perturbation
γ4 Perturbation
γ5 Perturbation
dL Perturbation
Cell Count
Time (Days)
10^6
10^4
10^2
10^0
0
200
400
600
S
P
S^L
P^L
TD_l
TD_m
TD_l^L
TD_m^L

D

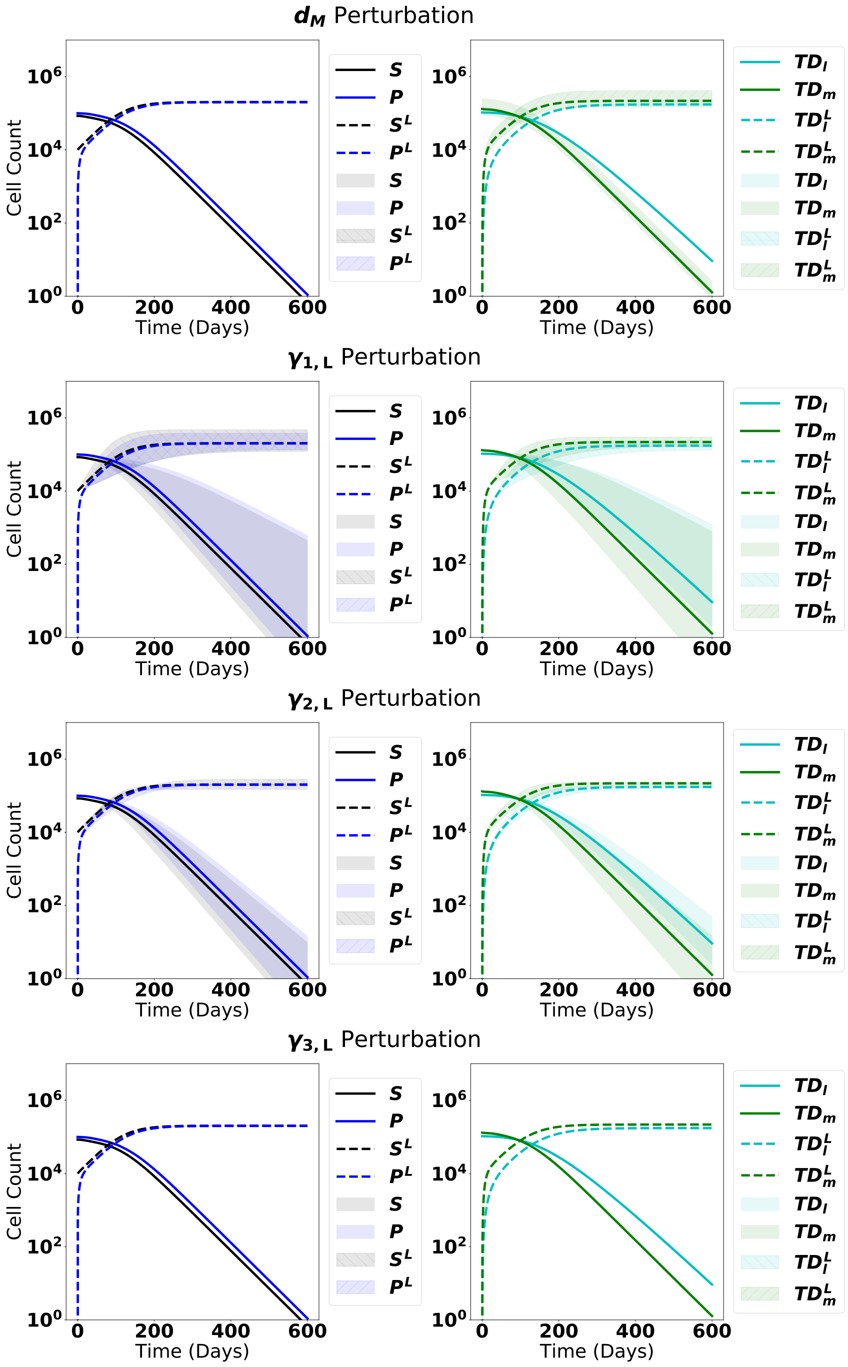

$d_M$ Perturbation
$\gamma_{1,L}$ Perturbation
$\gamma_{2,L}$ Perturbation
$\gamma_{3,L}$ Perturbation
Cell Count
Time (Days)
$10^0$
$10^2$
$10^4$
$10^6$
0
200
400
600
$S$
$P$
$S^L$
$P^L$
$TD_l$
$TD_m$
$TD_l^L$
$TD_m^L$

**E**

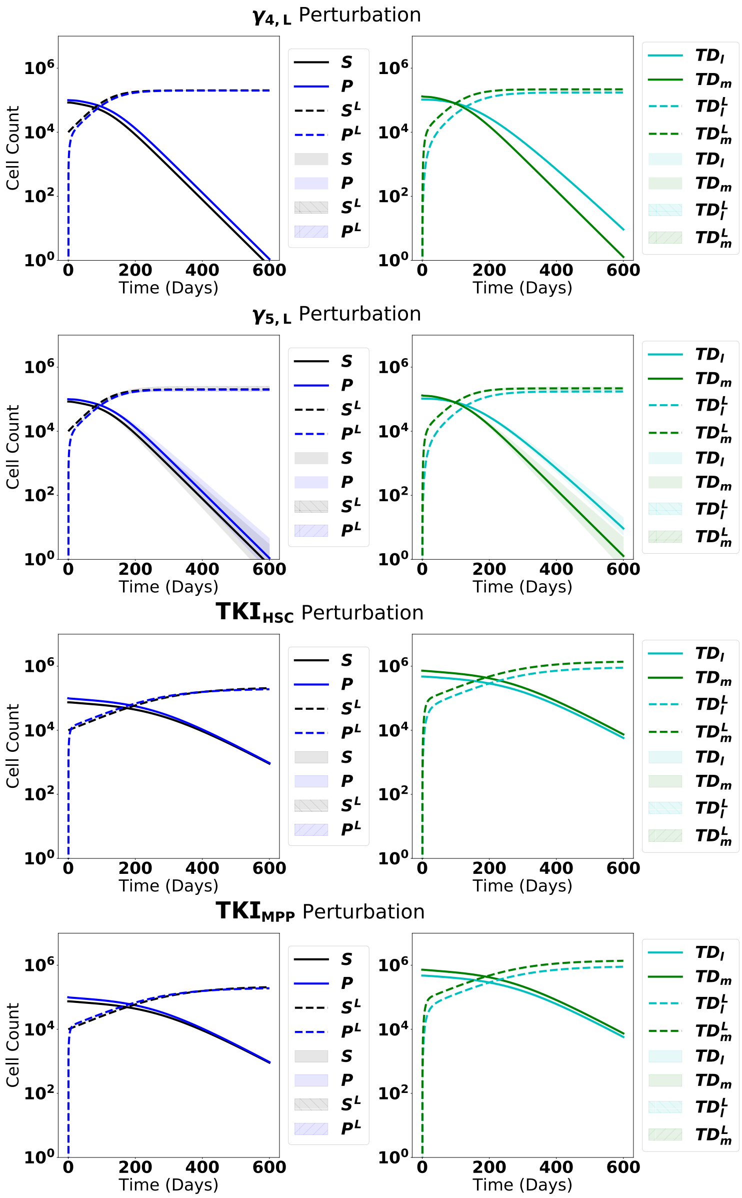


Figure S7: A-E: The dynamics from steady state upon introduction of leukemic stem cells with perturbations of each parameter, as labeled, ranging from 90% to 110% of the original parameter value. The solid line represents the median of the perturbations, while the shaded region represents 95% of the perturbation dynamics.

### 5 Simulations of transplant experiment

As described in the main text, we simulated a transplant experiment in a transgenic mouse model of CML performed in [5]. In this experiment, either leukemic stem $HSC^L$ or leukemic MPP $MPP^L$ cells were implanted into sub-lethally irradiated mice. Transplantation of $HSC^L$ enables engraftment and myeloid cell production that leads to CML. On the other hand, transplanting $MPP^L$ cells does not allow for long-term engraftment but results in a larger fraction of donor-derived lymphoid cells after 35 days We modeled this experiment by reducing the number of cells in equilibrium to mimic the effects of sublethal radiation (see Methods). Here, we present results of a range of possible reductions of $HSC^L$ and $MPP^L$ cells, and tracked the outcomes when 4000 $HSC^L$ or $MPP^L$ were introduced after the decrements from equilibrium. We then determine which of our parameter sets are consistent with the experimental outcomes found in [5]. The results are summarized in Figs. S8 and S9.

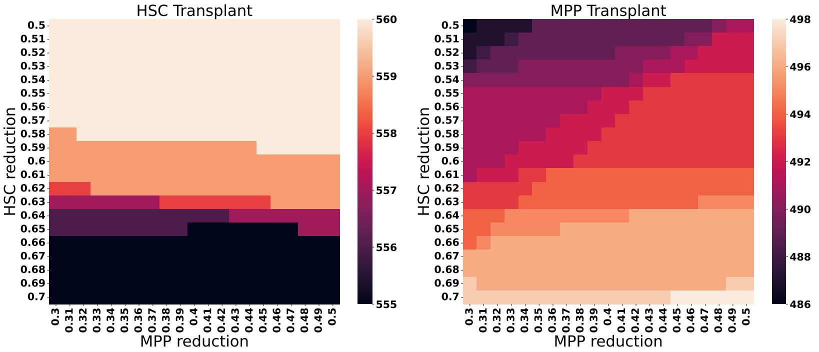


Figure S8: Heatmap depicting the outcomes of transplant experiments in the presence of decrements of 50-70% $HSC^L$ and 30-50% $MPP^L$ from their equilibrium values (see text for details). The heat map quantifies the number of parameter sets that are consistent with the experimental observations [5] that myeloid or lymphoid cells dominate after 35 days when $HSC^L$ or $MPP^L$ are transplanted, respectively.

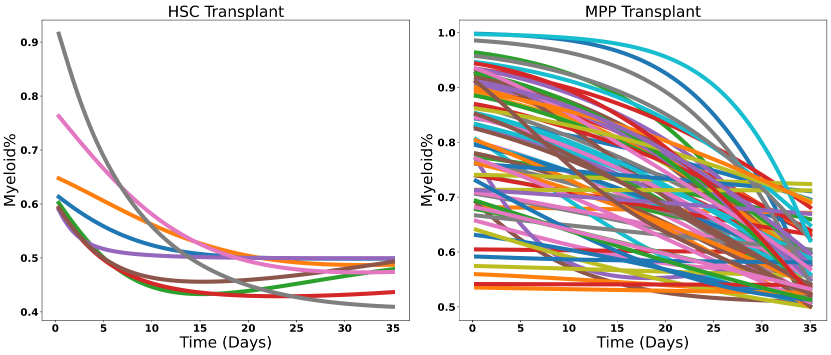


Figure S9: The dynamics of each parameter set that does not match the experimentally observed behavior of the transplant experiments from [5].These parameter sets are removed from the pool of eligible parameter sets.

The pairwise parameter distributions of the 478 remaining parameter sets are shown in Fig. S10.

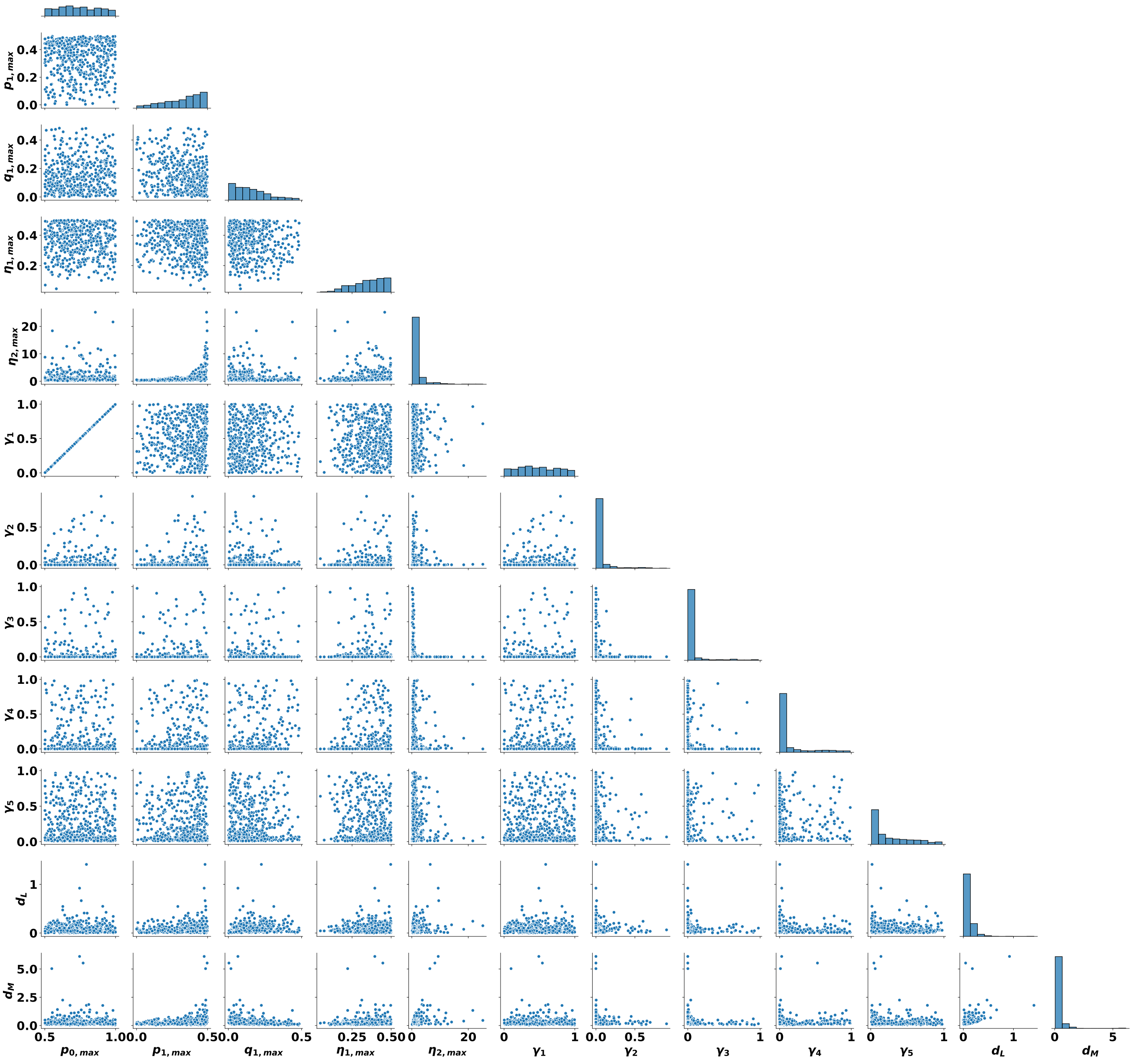


Figure S10: 478 parameters after removal determined through the depletion sweep.

### 6 Effective parameters

Here, we present the effective proliferation rates and self-renewal and branching factors–that is the values of these parameters that takes the feedback regulation into account. That is, the effective stem cell proliferation rate $\eta_1 = \eta_{1,max}/(1 + \gamma_2 x_S)$ in normal hematopoiesis and $\eta_1 = \eta_{1,max}/(1 + \gamma_2(x_S + x_{S^L}))$ when CML stem cells are present. The other effective parameters are defined analogously.

In Fig. S11, the effective HSC and MPP proliferation rates ($\eta_1$, $\eta_2$), the effective HSC and MPP self-renewal fractions ($p_0$, $p_1$) and branching fraction $q_1$ are shown. The corresponding effective parameters are shown in Fig. S12 when CML cells are introduced when the normal hematopoietic model system is at steady state and in response to treatment by TKIs (starts at the time labeled $t = 0$, as indicated by the vertical line) for the cases shown in Fig. 5 in the main text. In Fig. S13, we plot the effective proliferation, self-renewal and branching parameters for 50 parameter sets under treatment at early times.

### 7 Distributions of model parameters grouped by response to TKI treatment using synthetic data (478 parameter sets)

In Fig. S14, we plot the distributions of all the unregulated proliferation, self-renewal, and branching parameters, as well as the feedback gains, grouped by response to TKI therapy. In particular, the blue color indicates achievement of MR3 by 50 months (termed as responders) while the orange indicates that MR3 is not achieved within 50 months (termed as non-responders). The only parameters that clearly delineate the responders from non-responders are $p_{0,max}$ and $\gamma_1$, with responders occurring at the lower values and non-responders at the higher values.

### 8 Comparisons of prognostic criteria for predicting response to TKI therapy

#### 8.1 Performance of prognostic criteria using synthetic data (478 parameter sets)

Using data generated from our 478 parameter sets, we tested whether prognostic criteria could correctly identify patients who achieve MR3 within 18 months after therapy starts. We tested the performance of our prognostic criterion (relative change of transcript levels) against two existing clinical prognostics: halving time (the time it takes for the BCR-ABL1 transcripts to reach one-half of their pre-treatment value) and early molecular response (EMR) in which the fraction of BCR-ABL1 transcripts are 10% or less after 3 months of treatment. We also tested a prognostic criterion based on the ratio of transcript levels. These different prognostic criteria were calculated for both the first and second three months after the start of therapy (e.g., 0-3 months and 3-6 months). We calculated the corresponding receiving operating characteristic (ROC) curves and found the optimal threshold by maximizing the difference between true and false positive rates. The results are presented in Fig. S15 and reveal that generally the ratio and relative change prognostics offer similar performance, but that both demonstrate somewhat better performance compared to the traditional prognostics. In addition, all the prognostic criteria are more accurate when applied 3-6 months after the start of therapy than when applied during the 0-3 month period. We did not calculate the ROC curves for EMR but rather we only plotted the point that

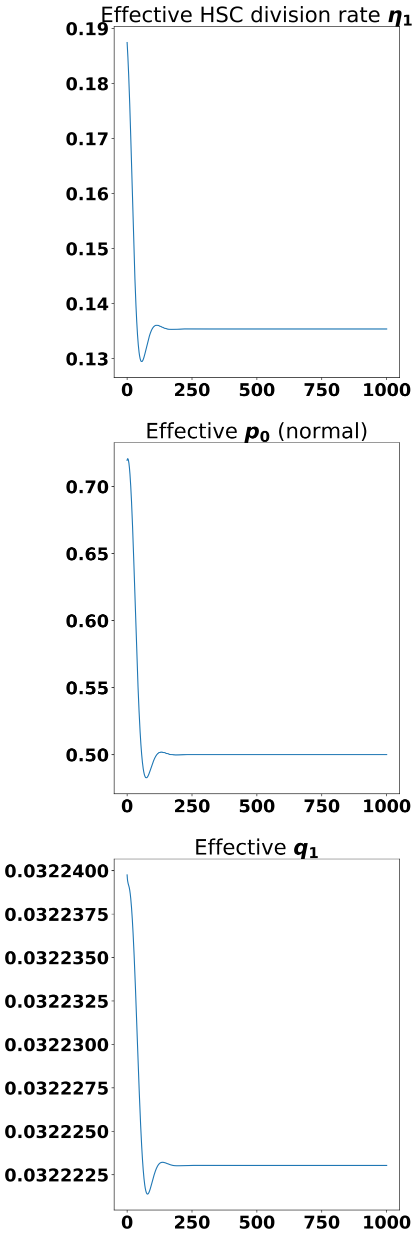


Figure S11: Effective parameters for proliferation, self-renewal and branching as the solutions to the model for the normal hematopoietic system approach steady state.

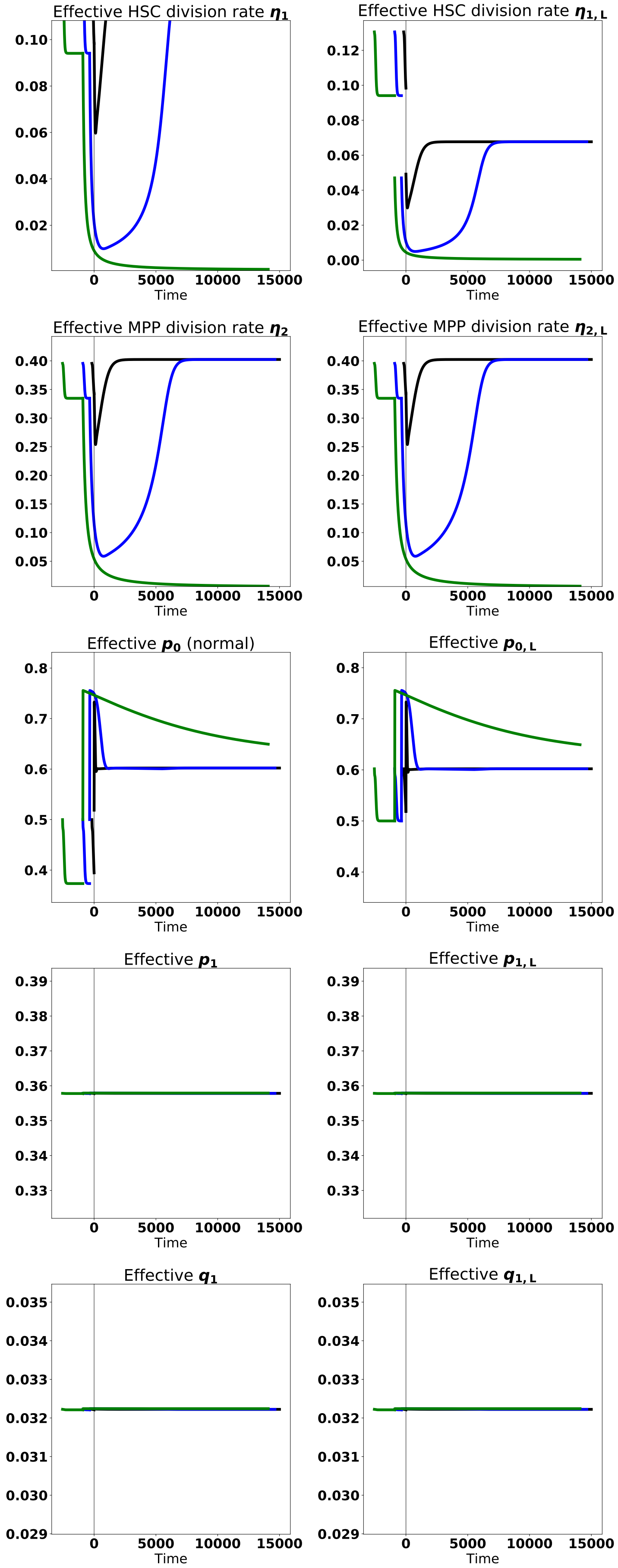


Figure S12: Effective parameters for proliferation, self-renewal and branching after leukemic stem cells are added to the normal system at steady state, and after TKI therapy begins for the cases shown in Fig. 5 in the main text. The vertical gray line depicts the start of treatment (time $t = 0$). Black represents treatment at early times (shortly after CML develops), blue– treatment at intermediate times, and green– treatment a long time after CML develops.

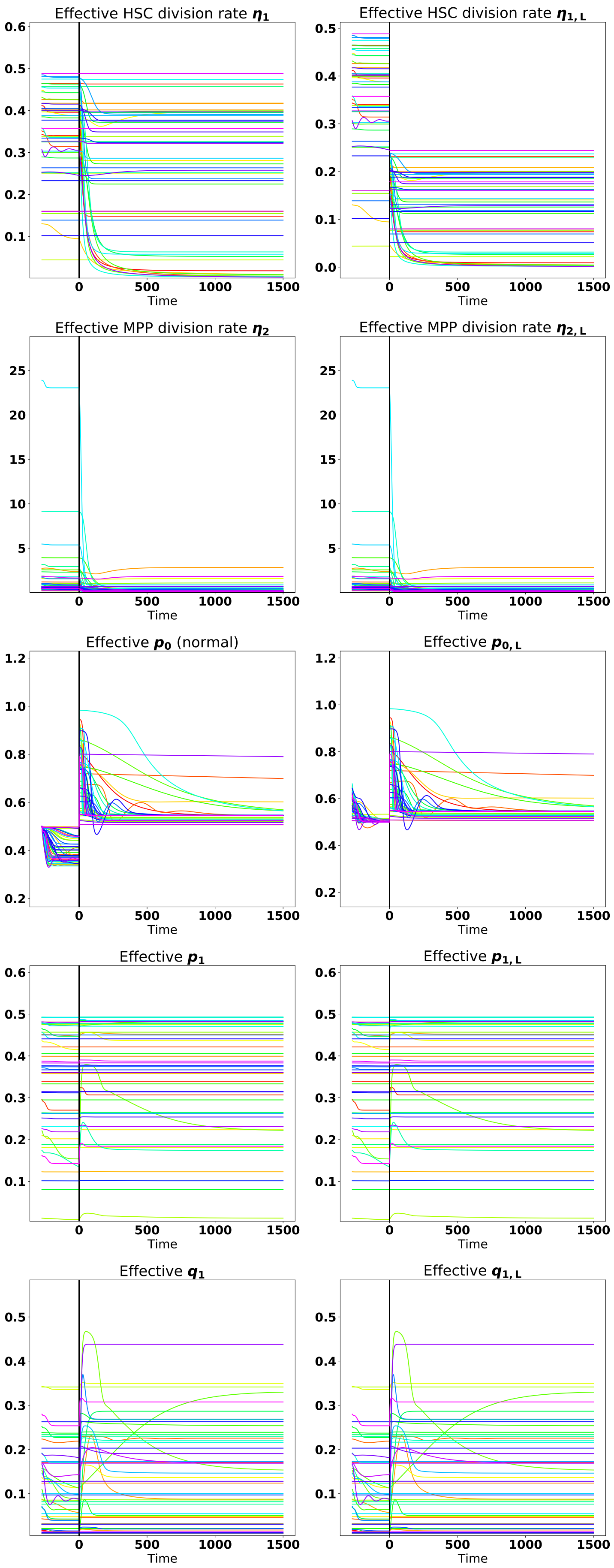


Figure S13: A spaghetti plot showing the dynamics of the effective proliferation, self-renewal and branching parameters for 50 parameter sets under TKI treatment started at early times.

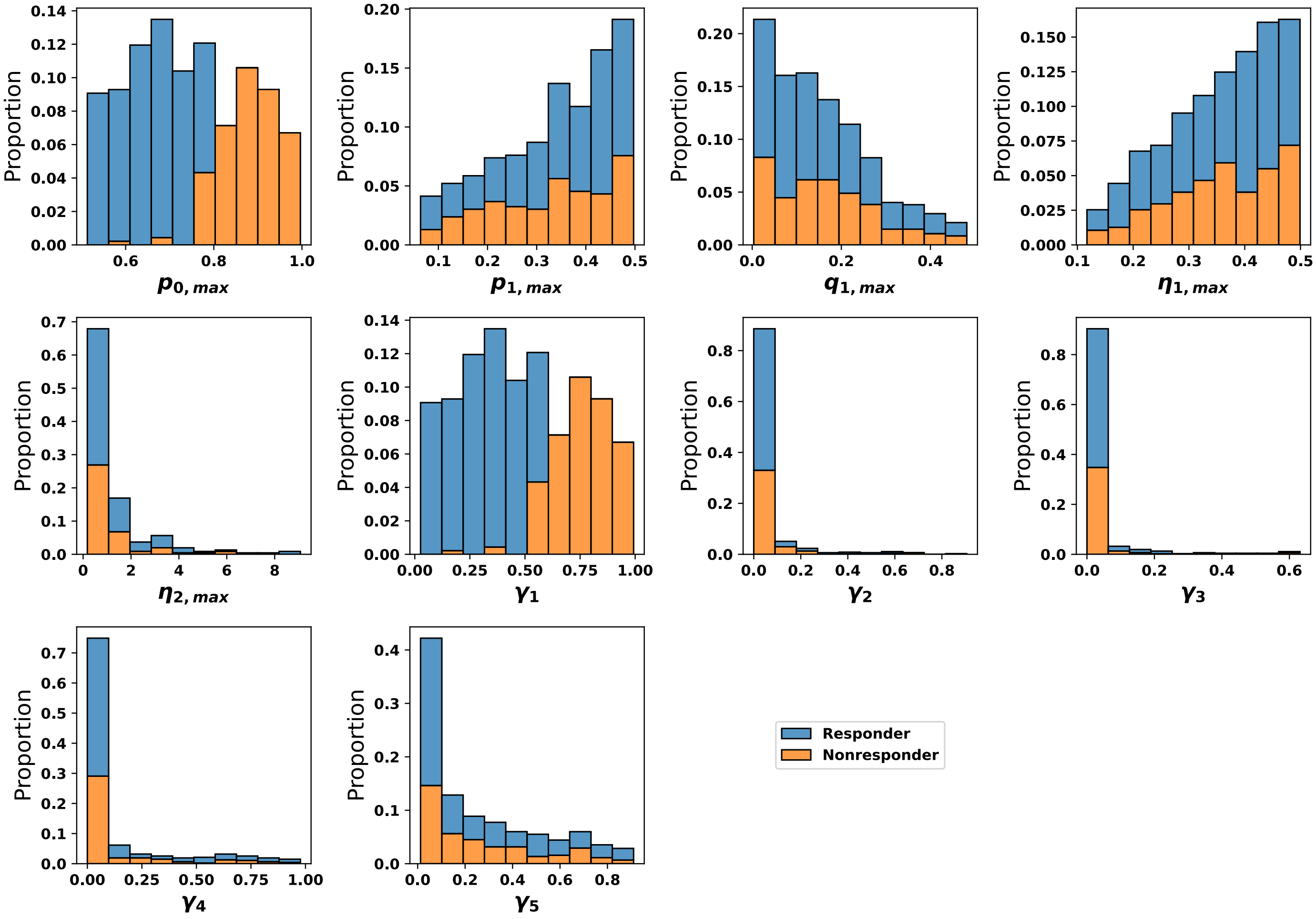


Figure S14: Parameter distributions by response to TKI treatment using synthetic data. Blue denotes a parameter set achieves MR3 within 50 months (responder) and orange denotes a parameter set that does not (non-responder).

corresponds to 10% transcript levels at 3 months (black circle) and 1% at 6 months (open black circle). The EMR prognostic criterion has fewer false positives but also fewer true positives than the other prognostics.

#### 8.2 Performance of prognostic criteria using patient data

The prognostic criteria were tested on anonymized patient data obtained from Dr. Van Etten's clinical practice, again asking whether MR3 at 18 months after treatment could be correctly predicted. We used data in which patients were kept on the same therapy for 6 months, either from the start of therapy, or after a change of therapy. For patients that achieved MR3 within 3 months, we did not include their data at 6 months. In Fig. S16, the first two columns correspond to results when the same therapy is applied for 6 months after patient diagnoses. The last two columns correspond to patients who have had

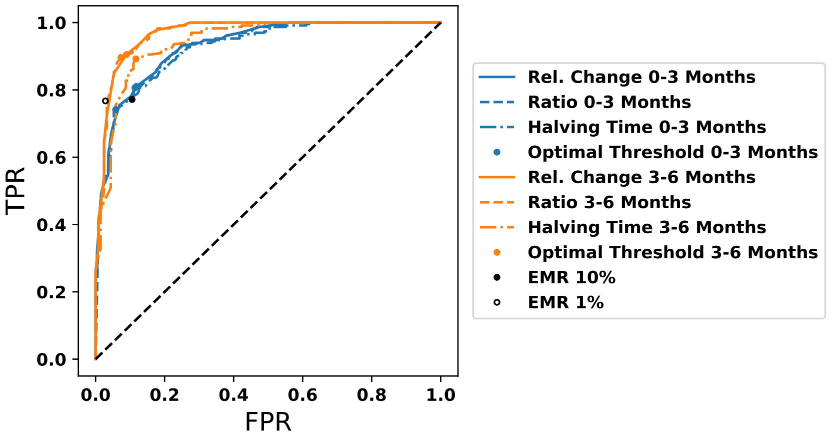


Figure S15: Comparison between our prognostic and alternate prognostics at the first and second three months after the start of therapy.

a change of therapy, but the new therapy is maintained for 6 months. We do not use the EMR as a prognostic in the cases when therapy is changed. The figure demonstrates that the prognostics are more accurate for the 3-6 month period, as predicted from the synthetic data. Although the numbers of patients are small, the relative ratio prognostic criterion is at least as accurate as, or more accurate than, the other criteria. In Fig. S17, the predictions of the prognostic criteria are grouped by whether the patients achieve or do not achieve MR3 by 18 months and by time period. In Fig. S18, the prognostic criteria data are aggregated into 3-month windows, where 0-3 months contains both 0-3 months after start of therapy and after a therapy change. The 3-6 months data is similarly aggregated. It is clear that the predictions using the 3-6 month data are more accurate than those using the 0-3 month window.

### 9 Combination therapy parameter distributions by response

Finally, in Fig. S19, we show how the distributions of model parameters from Fig. S15 change when TKI therapy is combined with a differentiation promoter. The blue hatching indicates a non-responder that becomes a responder while the orange hatching indicates that a responder becomes a non-responder. Here, response is defined as achieving MR3 in 50 months. We observe that differentiation therapy is very effective in driving non-responders at large $p_{0,max}$ and $\gamma_1$ to become responders, but also drives parameter sets that responded to TKI monotherapy at small values of $p_{0,max}$ and $\gamma_1$ to no longer achieve MR3 at 50 months (non-responders).

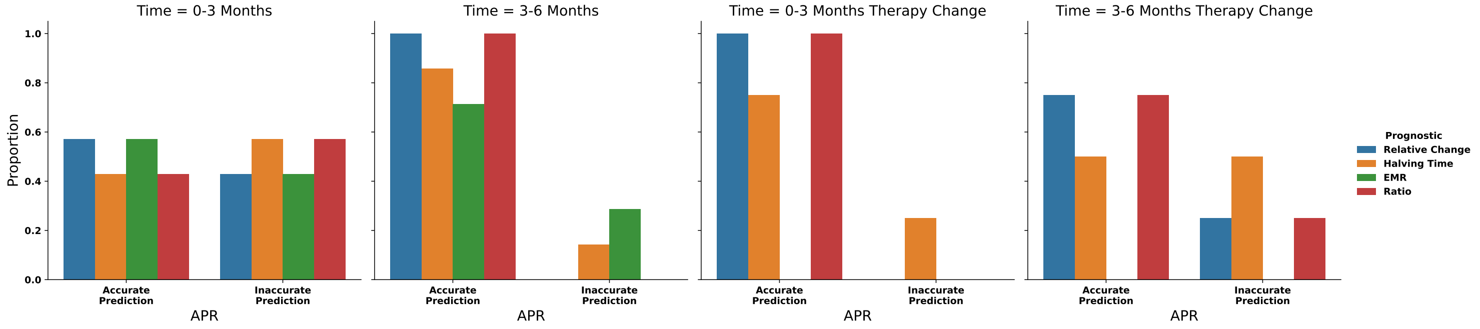


Figure S16: The ability of prognostic criteria to predict MR3 by 18 months is evaluated using anonymized patient data in which patients received the same therapy for 6 months, either from the start of therapy or after a change in therapy. The first two columns correspond to results when the same therapy is applied for 6 months after patient diagnoses. The last two columns correspond to patients who have had a change of therapy, but the new therapy is maintained for 6 months. The EMR prognostic criterion is not used when the patients have had a therapy change.

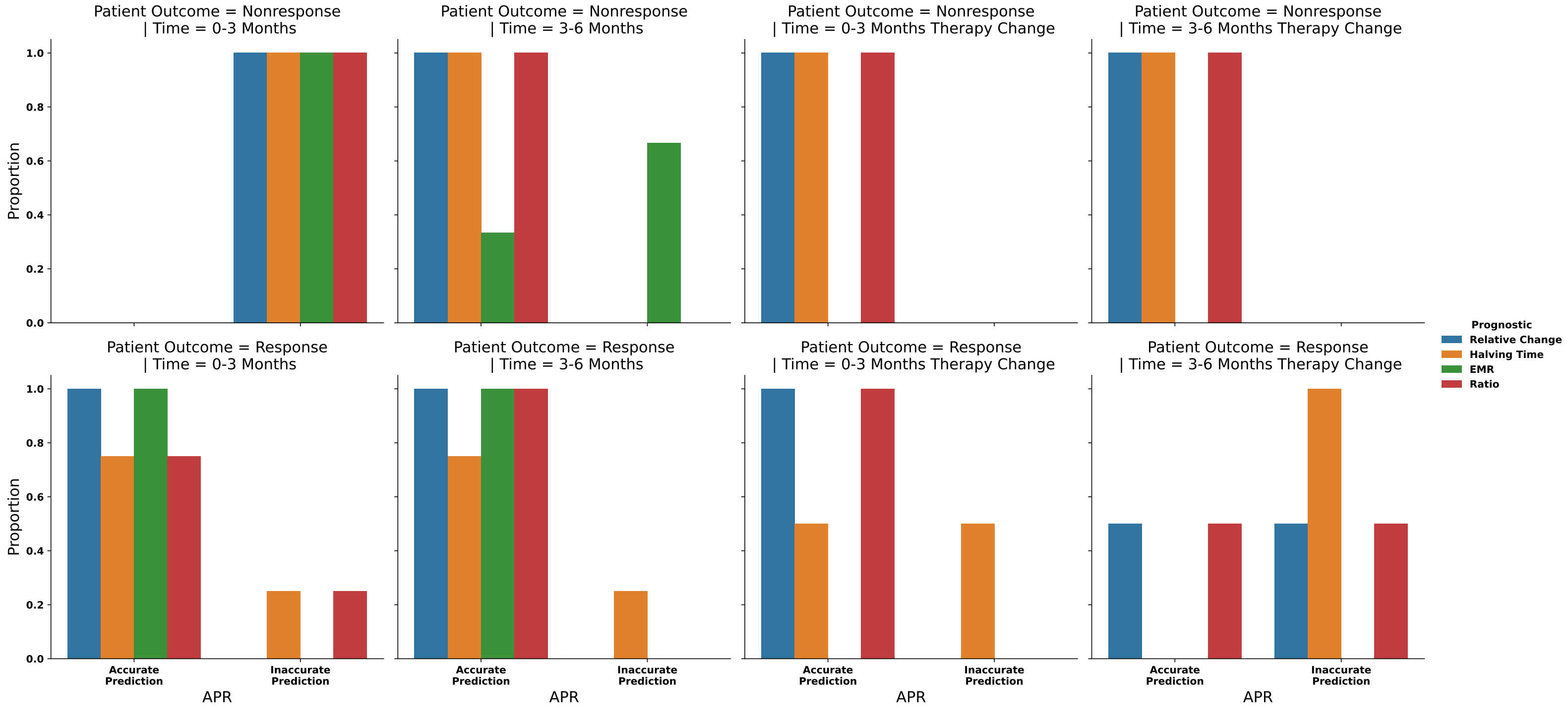


Figure S17: The predictive ability of each prognostic criterion from Fig. S16 but grouped based upon patient outcome (blue– responder and orange– nonresponder) and the time frame for prediction.

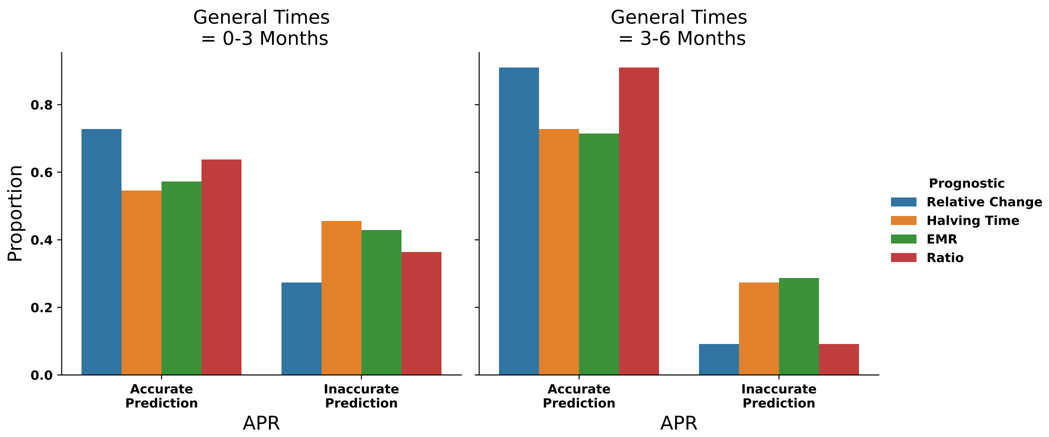


Figure S18: Aggregating the accuracy of prognostic criteria predictions from Fig. S16 where a general time of 0-3 months contains both 0-3 months after start of therapy and after therapy change. The 3-6 month data is aggregated similarly.

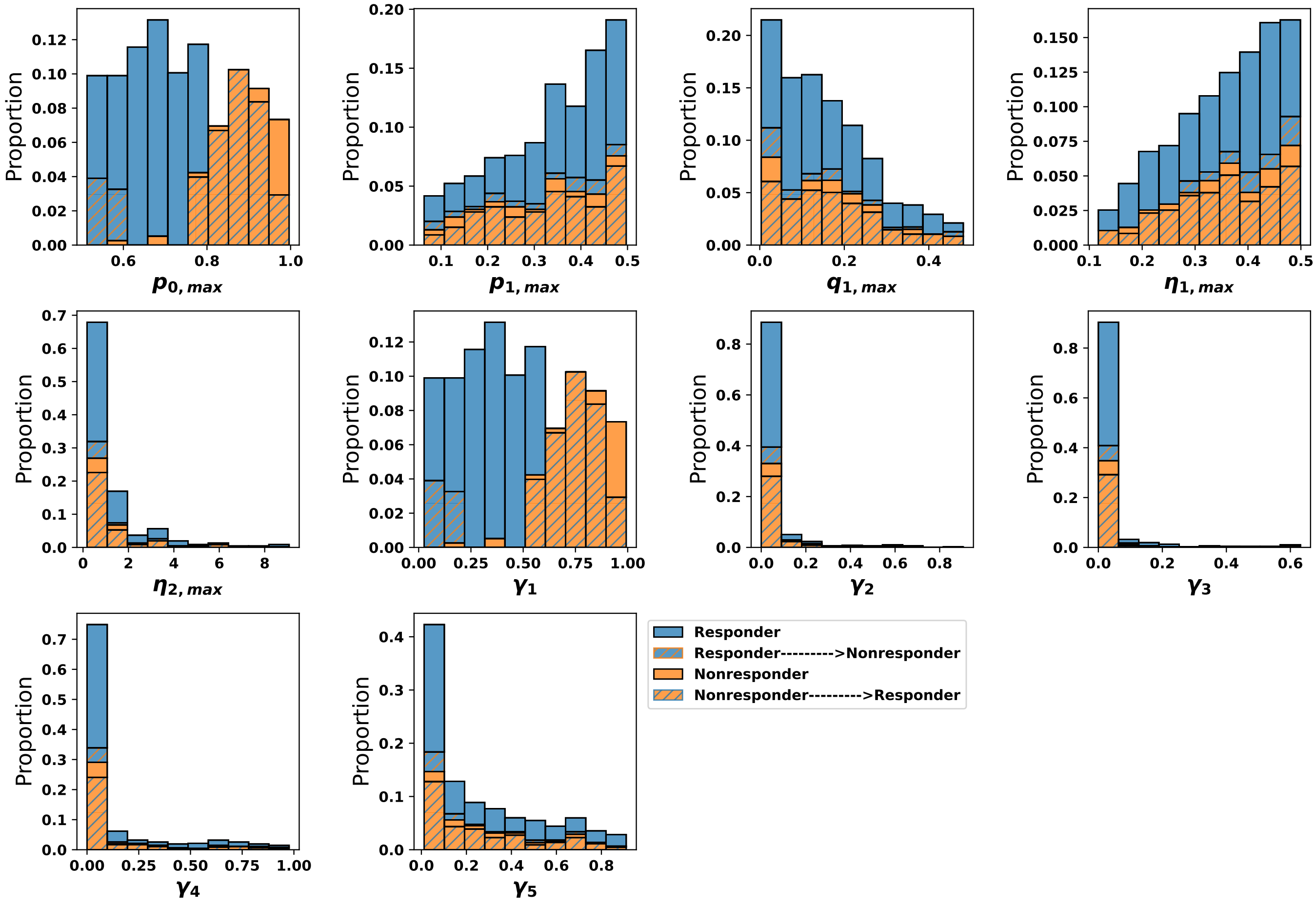


Figure S19: Changes in the parameter distributions from Fig. S14 when combined differentiation and TKI therapy is administered. Blue denotes a parameter set achieves MR3 within 50 months and orange denotes parameter sets that do not. Orange hatching denotes a parameter set that achieves MR3 within 50 months with TKI monotherapy, but does not achieve MR3 within 50 months under combination therapy. Blue hatching denotes a parameter set that does achieve MR3 within 50 months with combination therapy but does not with TKI therapy alone.
